# Whole body regeneration deploys a rewired embryonic gene regulatory network logic

**DOI:** 10.1101/658930

**Authors:** Hereroa Johnston, Jacob F. Warner, Aldine R. Amiel, K Nedoncelle, João E Carvalho, Eric Röttinger

**Affiliations:** Université Côte d’Azur, CNRS, INSERM, Institute for Research on Cancer and Aging, Nice (IRCAN), Nice, France; Université Côte d’Azur, Institut Fédératif de Recherche – Ressources Marines, (MARRES), Nice, France

## Abstract

For over a century, researchers have been trying to understand the relationship between embryogenesis and regeneration. A long-standing hypothesis is that biological processes implicated in embryonic development are re-deployed during regeneration. In the past decade, we have begun to understand the relationships of genes and their organization into gene regulatory networks (GRN) driving embryonic development and regeneration in diverse taxa.

Here, we compare embryonic and regeneration GRNs in the same species to investigate how regeneration re-uses genetic interactions originally set aside for embryonic development. Using a well-suited embryonic development and whole-body regeneration model, the sea anemone *Nematostella vectensis*, we show that at the transcriptomic level the regenerative program partially re-uses elements of the embryonic gene network along with a small cohort of genes that are specifically activated during the process of regeneration. We further identified co-expression modules that are either i) highly conserved between these two developmental trajectories and involved in core biological processes (e.g., terminal differentiation) or ii) regeneration specific modules that drive cellular events, such as apoptosis, that are unique to regeneration.

Our global transcriptomic approach suggested that regeneration reactivates embryonic gene modules following regeneration-specific network logic. We thus verified this observation by functionally dissecting the role of MEK/ERK signaling during regeneration and established a first blueprint of the regeneration MEK/ERK-dependent GRN in *Nematostella*. Comparing the latter to the existing GRN underlying embryogenic development of the same species, we show at the network level that i) regeneration is a partial redeployment of the embryonic GRN, ii) embryonic gene modules are rewired during regeneration and iii) they are interconnected to novel down-stream targets, including “regeneration-specific” genes.

**Significance statement:** In this intra-species transcriptomic comparison of embryonic development and regeneration in a whole-body regeneration model, the sea anemone *Nematostella vectensis*, we identified that 1) regeneration is a transcriptionally modest event compared to embryonic development and 2) that although regeneration re-uses embryonic genetic interactions, it does so by using regeneration specific network logic. In addition to identifying that apoptosis is a regeneration-specific event in *Nematostella*, this study reveals that GRN modules are reshuffled from one developmental trajectory to the other, even when accomplishing the same task (*e.g.* forming a fully functional organism). These findings highlight the plasticity of network architecture and set the basis for determining and functionally dissecting regeneration-inducing regulatory elements. From an evolutionary perspective, our study sets the foundation for further comparative work and provides new opportunities to understand why certain organisms can regenerate while others cannot.

## Introduction

Regeneration of cells, tissues, appendages or even entire body parts is a widespread yet still rather poorly understood phenomenon in the animal kingdom ^1^. A long-standing question in the field of regeneration is whether and to what extent embryonic gene programs, initially used to build an organism, are re-used during regeneration ^2^. Several transcriptomic studies of regeneration in axolotl, anole, zebrafish and sea anemones have highlighted the importance of the re-deployment of developmental pathways ^3–8^. Many studies have directly compared embryonic and regenerative gene expression focusing on single or groups of genes, thus identifying i) genes that are specific to embryonic development (Binari et al. 2013), ii) genes that are specifically expressed or required during regeneration ^9,10^, and iii) embryonic genes that are re-used during regeneration to some extent ^11–15^. One recent study has addressed the question by comparing the transcriptional dynamics between post-larval development and regeneration from dissociated cells in sponges, further highlighting partial similarities between these two developmental trajectories ^16^. Another recent study has investigated the relationship between larval skeleton development and brittle star arm regeneration with a special emphasis on FGF signaling, revealing that a skeleton developmental gene regulatory module is re-deployed during regeneration ^17^. Investigating the transcriptomic relationship between leg development and regeneration in a marine arthropod, the authors describe that both processes involve a similar set of genes, although their temporal relationship appears to be different ^18^. To date, however, no large-scale functional comparison between embryonic development and whole-body regeneration within the same organism has been carried out.

The sea anemone *Nematostella vectensis* (Cnidaria, Anthozoa) is an embryonic development and whole-body regeneration model that is ideally suited for this line of inquiry (Fig. 1A, ^19^). *Nematostella* has long been used as a model system to study embryonic development, evolution of body patterning, and gene regulatory networks ^20–25^. More recently, *Nematostella* has emerged as a powerful whole-body regeneration model as it is capable of re-growing missing body parts in less than a week ^8, 26–34,32^. Regeneration in *Nematostella* follows a dynamic but highly stereotypical morphological and cellular program involving tissue re-modeling and the *de novo* formation of body structures (Amiel et al. 2015). Initiation of regeneration requires a crosstalk between tissues and two populations of fast and slow cycling, potential stem cell populations ^32^. While many developmental signaling pathways are deployed during regeneration ^8,28,33,34^, their regulatory logic remains unknown.

**Figure 1:**
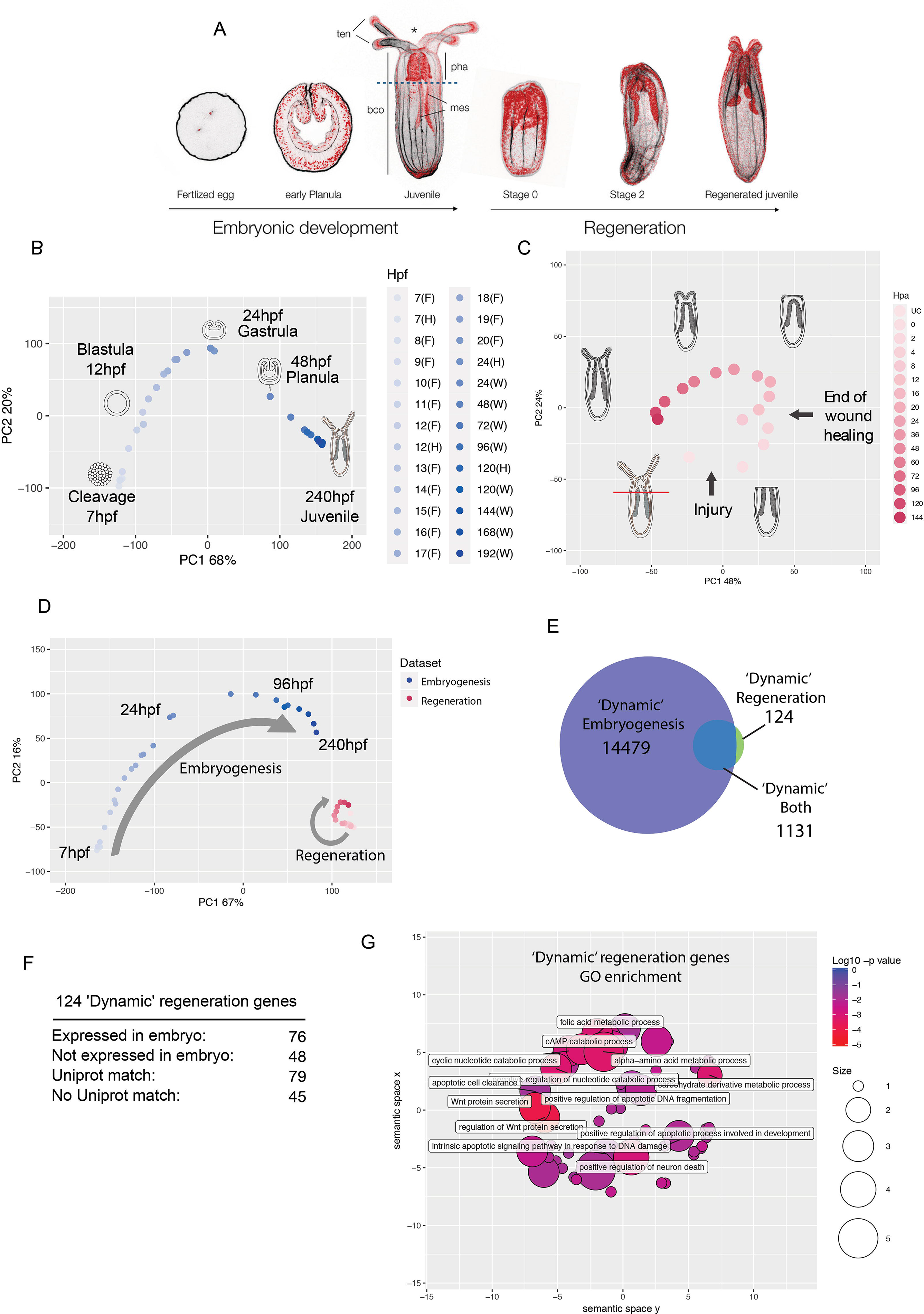
Comparison of embryonic and regenerative transcriptomes. (**A**) General morphology of *Nematostella* during embryonic development and regeneration (black; f-actin/Phalloidin, red; nuclei/DAPI). **(B)** Principal component analysis (PCA) of three embryonic datasets: ^34–36^. Plot legend indicates timepoint and dataset (Fischer et al.: F, Helm et al.:(H) and Warner et al.:(W). The majority of variation is observed in the first 24 hours of development. (**C**) PCA of regeneration dataset sampled at Uncut (UC), 0, 2, 4, 8, 12, 16, 20, 24, 36, 48, 60, 72, 96, 120, 144 hpa. Regeneration proceeds through a wound-healing phase (0-8 hpa) followed by the early regenerative program (12-36 hpa) and ending with a late regenerative program which approaches the uncut condition (48-144 hpa). (**D**) PCA of embryonic versus regeneration samples. Embryogenesis (blue) exhibits far greater transcriptomic variation than regeneration (red). (**E**) Comparison of differentially expressed (|log_2_(FC)| >2 & FDR < 0.05 for any timepoint comparison against t_0_ where t_0_ = 7hpf for embryogenesis and 0hpa for regeneration) ‘dynamic’ genes during embryogenesis (blue) and regeneration (green). (**F**) Details of the regeneration specific genes expression and classification. (**G**) GO term enrichment for dynamic regeneration genes.

Here, we take advantage of *Nematostella* to address the historical question to what extent regeneration recapitulates embryonic development and to decipher molecular signatures unique to regeneration. In the present study, we performed a global transcriptomic comparison of embryogenesis and regeneration using deeply sampled transcriptomic datasets. This approach was combined with a pathway-specific functional analysis as well as a comparison of the embryonic and regeneration GRNs within the same species. Overall, this study i) revealed that whole-body regeneration is transcriptionally modest in comparison to embryonic development; ii) identified genes, cellular processes and network modules specific to the process of regeneration; iii) demonstrated the plasticity of the network architecture underlying two developmental trajectories; iv) and showed that whole-body regeneration deploys a rewired embryonic GRN logic to reform lost body parts.

## Results

### Regeneration is transcriptionally modest compared to embryonic development

To compare embryogenesis and regeneration on a global transcriptome-wide scale we employed four RNAseq datasets, one spanning 16 time points of regeneration ^34^ and three spanning a total of 29 embryonic time points ^34–36^. In order to directly compare the data, raw sequencing reads were processed, mapped and quantified using the same workflow for all datasets (see materials and methods for quantification details). As the embryonic data were the result of several previous studies we applied a batch correction ^37^ using developmental time-point as a categorical covariate (Fig. S1). To assess the transcriptomic states underlying embryogenesis we performed principal component analysis (PCA) on batch corrected embryonic data (Fig. 1B). We found that the majority of gene expression changes occur during the first day of embryonic development from cleavage to blastula stage (Fig. 1B, 7 hours post fertilization (hpf) – 24hpf, PC1 proportion of variance 68%; PC2 proportion of variance 20%) indicating large transcriptomic differences in early embryogenesis. From 96hpf onwards the samples exhibited modest changes in transcriptional variation suggesting that the majority of transcriptional dynamics are driving embryogenesis are complete by this stage (96hpf-240hpf).

When we examined the regenerative program using PCA (Fig. 1C), we observe four distinct transcriptional programs: i) a wound-healing phase (0-8hpa) that is followed by ii) the activation of the early regenerative program (8-20 hours post amputation (hpa) in which the samples are distributed along the second principal component (PC2 proportion of variance = 24%, Fig. 1C). iii) From 20hpa onwards, the majority of variation in gene expression is explained by the first principal component during the late regenerative phase (PC1 proportion of variance = 48%, 20hpa-144hpa, Fig.1C). iv) Towards the end of regeneration (144hpf), we observe a transcriptomic profile approaching the uncut samples indicating a return to steady state. These profiles correlate with the major events of oral regeneration in *Nematostella* and indicate that our sampling strategy effectively covers the major transcriptional hallmarks of regeneration ^31^.

We next directly compared the transcriptomic variation of regeneration and embryogenesis using PCA and found that the transcriptional changes during regeneration were relatively modest compared to those observed during embryogenesis with the vast majority of variation in the first two principal components being driven by the embryonic data (PC1 proportion of variance = 67%, PC2 proportion of variance = 16%, Fig 1D). This indicates that the transcriptional dynamics of embryogenesis are more profound than those of regeneration. This finding was buttressed by comparing the number of ‘dynamically expressed genes’, those which are significantly differentially expressed (|log_2_(fold change)|>2 and false discovery rate < 0.05) at any time point compared to t_0_ defined as 0hpa for regeneration and 7hpf (the onset of zygotic transcription) for embryogenesis. Embryogenesis exhibited more than ten times the number of dynamically expressed genes compared to regeneration (15610 and 1255 genes respectively, Fig. 1E). These results show that regeneration, when compared to embryogenesis, employs far fewer dynamic genes to accomplish a similar task: (re)constructing a functional animal. Of the dynamic genes observed during regeneration the vast majority (1131 out of 1255) are also dynamically expressed during embryogenesis, demonstrating that regeneration is largely a partial re-use of the embryonic gene complement (Fig. 1E).

### Identification of dynamically expressed genes specific to regeneration

Among the genes dynamically expressed during regeneration, a small fraction, 124 genes, exhibit differential expression only during regeneration which we term “regeneration specific” (Supplementary table 1). Furthermore, 48 of these genes are detectable specifically during the regeneration process indicating they are transcriptionally silent during embryogenesis (Fig. 1F). The remaining 76 genes are expressed during embryonic development but are not considered dynamic from our differential expression analysis above (Fig. 1F). Interestingly, several of the 124 “regeneration specific” genes, for example *wntless* (jgi|Nemve1|100430) and *agrin* (jgi|Nemve1|196727), have previously been reported to be important regulators of regeneration in bilaterians ^38,39^. Furthermore, among the “regeneration specific” genes, 45 have no known homology in the Uniprot database (BLASTp, e-value cutoff <0.05, see methods for annotation details), suggesting to be possible taxonomically-restricted genes. These results indicate not only a possible evolutionary conservation of gene use in regeneration, but also identify additional genes that may play important roles specifically during whole body regeneration. A gene ontology (GO) term enrichment analysis on these 124 “regeneration specific” genes, revealed a suite of biological process GO terms relating to modulation of signaling pathways (e.g. *wntless*), metabolic processes and apoptotic cell death, indicating an essential role for these processes in regeneration (Fig. 1G).

### Embryonic gene modules are partially re-deployed during regeneration

In terms of dynamically expressed genes, regeneration uses less than one tenth the number of genes compared to embryonic development. We were thus interested in how these genes were deployed and arranged into co-expression networks. We sought to determine if embryonic gene network modules themselves are reused in a reduced capacity or if regeneration deploys novel gene module arrangements. To investigate this, we first used fuzzy c-means clustering to group the genes by expression profile ^40^. We regrouped the gene expression profiles into eight embryonic clusters (Fig. 2A) and nine regeneration clusters (Fig. 2B) ^34^. To explore these expression clusters (also named modules), we performed GO-term enrichment for each cluster (Table S1, Table S2) and examined the clusters at the gene level. We found that modules that were activated early in both processes (embryogenesis cluster 4, regeneration cluster 6, Fig. 2A,B) contained several canonical developmental genes: *wntA* (jgi|Nemve1|91822), *lmx* (jgi|Nemve1|95727) and *foxA* (jgi|Nemve1|165261) in embryonic cluster 4 (Fig. 2Ai) and *tcf* (jgi|Nemve1|132332), *spr* (jgi|Nemve1|29671), and *runx* (jgi|Nemve1|129231) in regeneration cluster 6 (Fig. 2Bi). The early embryonic transcriptional activation of *wntA, lmx and foxA* has been previously reported ^22,41–43^ and confirmed by *in situ* hybridization at 24hpf (Fig. 2Aii). The spatiotemporal expression patterns of *tcf*, *spr* and *runx* during regeneration (uncut, 0-60hpa, Fig. 2Bii) confirm the dynamic expression patterns of these genes as early as 2hpa at the amputation site (Fig. 2Bii c,c’, c’’). These data also suggest that these genes function as a synexpression group ^44^, first activated at the amputation site, then in the mesenteries as regeneration proceeds. Furthermore, we conclude from this analysis that classical developmental genes are involved in the early phases of both embryogenesis and regeneration.

**Figure 2:**
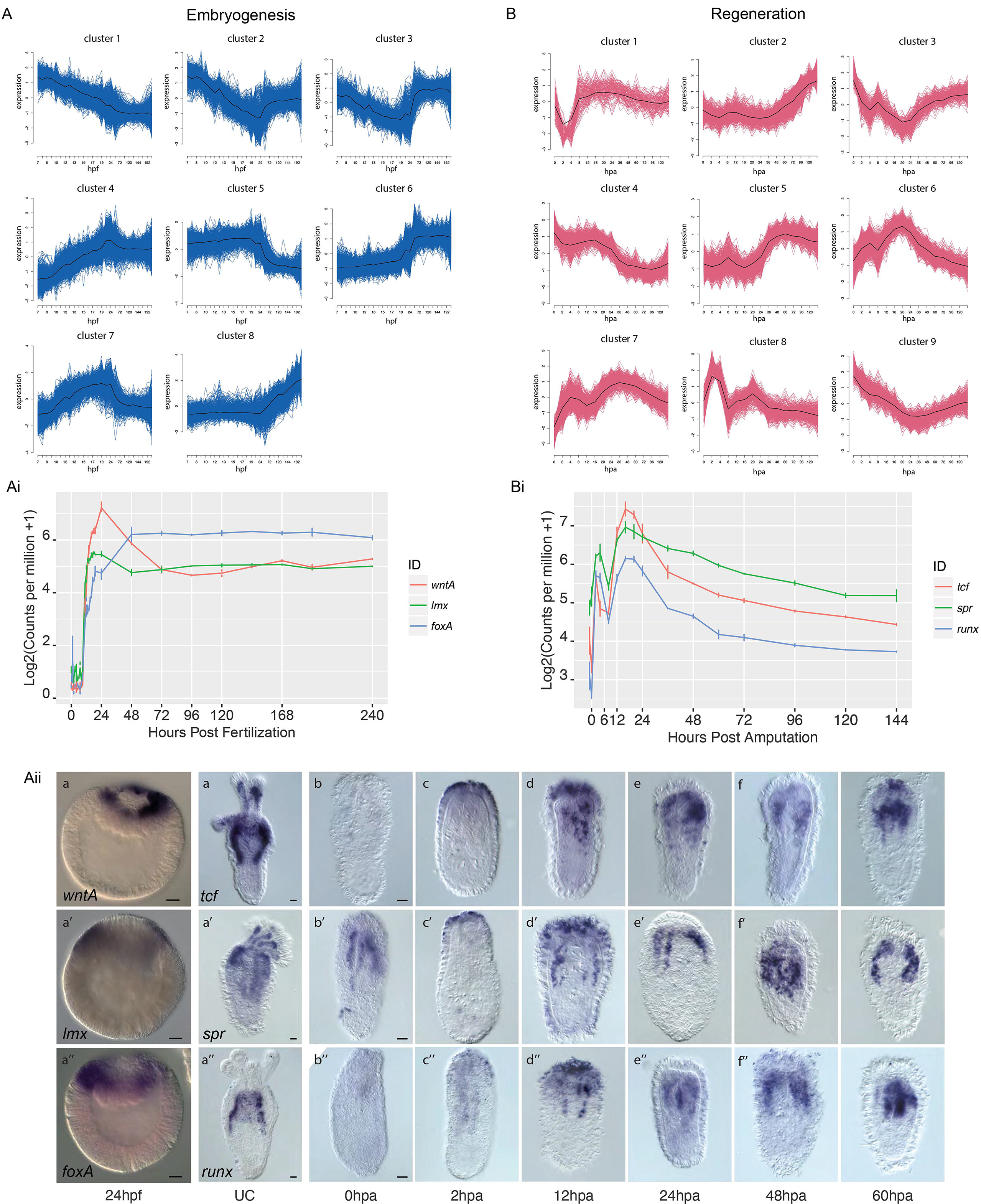
Embryonic and Regenerative gene expression forms discrete clusters. (**A, B**) Fuzzy c-means clustering of embryonic (**A**) and regeneration (**B**) gene expression. Each cluster is plotted with standardized expression along the y-axis and developmental time along the x-axis. Black trace denotes the cluster core (centroid). (**Ai-Aii**) Exemplar gene expression from a cluster activated early during embryogenesis in the cluster E-4. *wntA*, *lmx*, *foxA* are temporally co-expressed (**Ai**) and *in situ* hybridization at 24hpf confirms early activation of this gene cluster (**Aii**). (**Bi-Bii**) Exemplar gene expression from a cluster activated in the early regenerative program. *tcf*, *spr*, *runx* are all activated early in the cluster R-6 and are temporally co-expressed (**Bi)** and *in situ* hybridization during regeneration confirms early activation and the expression dynamics of this gene cluster (**Bii**). Scale bar: 20µm

We next analyzed whether the same groups of genes were co-regulated during embryogenesis and regeneration, by testing if gene expression observed during both processes were arranged in similar co-expression modules (Fig. 3). We compared regeneration and embryonic clusters on a gene-cluster membership basis to identify significant overlaps. Regeneration clusters with high overlap of a specific embryonic cluster indicate a shared or re-used network logic since the same suite of genes are deployed as a bloc in both processes. Regeneration clusters with low overlap to any single embryonic cluster on the other hand are likely to be *de novo* genetic arrangements specific to regeneration.

**Figure 3:**
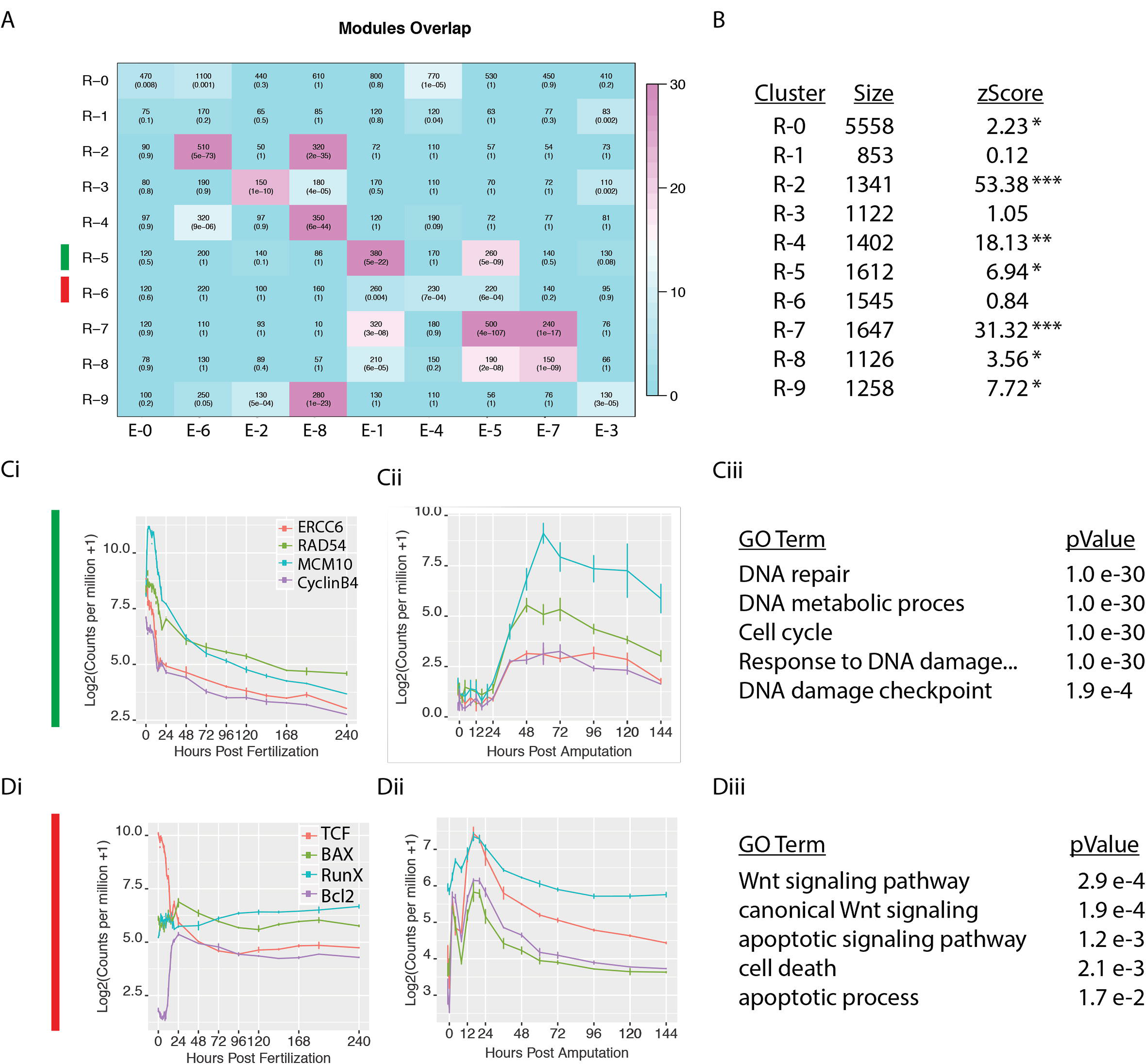
Embryonic gene modules are partially redeployed during regeneration. (**A**) Overlap table of regeneration versus embryonic modules. In each cell, the overlap itself is quantified along with the pvalue (fischers exact test). Color indicates –log_10_(pvalue), with a brighter magenta indicating a more significant overlap. R-0 and E-0 contain genes that are not assigned to any module. (**B**) Table indicating the size, and the co-clustering zStatistic. A zStatistic >2 (*) indicates moderate module conservation, >10 (**) high conservation > 30 (***) very high conservation). (**C**) The conserved module R-5 with exemplar genes (*ercc6, rad54B*, *mcm10, cyclinB3*) showing coexpression during embryogenesis (**Ci**) and regeneration (**Cii**). GO-term enrichment identifies terms associated with cell proliferation (**Ciii**). (**D**) The regeneration specific module R-6 with exemplar genes (*tcf*, *bax*, *runx*, and *bcl2*) showing co-expression during regeneration (**Di**) but divergent expression during embryogenesis (**Dii**). GO-term enrichment identifies terms associated with apoptosis and wnt signaling (**Diii**).

We found that the majority of the regeneration clusters exhibited significant overlap with one or more embryonic clusters (Fig. 3A, B). These ‘conserved modules’ also exhibited high preservation permutation co-clustering zStatistics (Fig 3B; >2 indicating conservation; >10 indicating high conservation; permutations = 1000) ^45^. Importantly, we identified two clusters, R-1 and R-6, which exhibited relatively low overlap with any one embryonic cluster, indicating that these are likely ‘regeneration specific’ arrangements. When we examined the GO-term enrichment of each cluster, we found that in general, highly conserved clusters (e.g. R-5) were enriched in GO-terms corresponding to homeostatic cell processes while lowly conserved regeneration specific clusters (e.g. R-6) were enriched in GO-terms describing developmental signaling pathways (Fig. 3A, Supplementary tables 2 & 3). These results suggest that the gene module arrangements pertaining to core biological functions such as cell proliferation are common to embryogenesis and regeneration while the gene modules containing developmental patterning genes are unique to each process.

Two clusters that exemplify these findings are R-5 and R-6. Cluster R-5, a conserved cluster (zStatistic 6.94) showed strong enrichment of cell-proliferation related GO-terms (Fig. 3Ciii). When we examined exemplar genes (with intra module membership scores >0.95) *ercc6-like* (jgi|Nemve1|110916), *rad54B* (jgi|Nemve1|209299), *mcm10* (jgi|Nemve1|131857), *cyclinB3* (jgi|Nemve1|208415), we observed co-expression patterns that correlate well to the timing of proliferation activation during *Nematostella* regeneration with an activation at 24hpa, a peak at 48hpa, and a taper off thereafter ^29,31^ (Fig. 3Cii). These exemplar genes are also co-expressed during embryogenesis (cluster E-1, Fig. 3Ci), further demonstrating module conservation. In contrast to cluster R-6 is specific to regeneration. This module exhibits strong enrichment of GO-terms relating to apoptosis and developmental signaling pathways. When we examined 4 exemplar genes *tcf* (jgi|Nemve1|132332), *bax* (jgi|Nemve1|100129), *runx* (jgi|Nemve1|129231), *bcl2* (jgi|Nemve1|215615), we observed co-expression during regeneration (Fig 3Dii) but divergent profiles during embryogenesis (Fig 3Di) indicating that this grouping of genes is indeed ‘regeneration specific’. These results suggest that modules containing genes responsible for basic cellular functions are largely re-used and co-expressed between embryogenesis and regeneration, while those including genes that are important for the activation of developmental processes have regeneration-specific arrangements.

### Apoptosis is specifically required for regeneration in Nematostella

Having observed a strong enrichment for apoptosis related GO-terms in the list of 124 regeneration-specific genes (Fig. 1G) and in the regeneration specific module R-6 (Fig. 3Dii-Diii), we investigated the role of apoptosis during the regenerative process. Several genes relating to apoptosis, including the regeneration-specific genes *bax* (jgi|Nemve1|100129), *caspase-3* (jgi|Nemve1|100451), *bcl2* (jgi|Nemve1|215615), and an additional *bcl2* (which we term *bcl2B*, jgi|Nemve1|128814), belong to module R-6 and are activated shortly after amputation (Fig. 4Ai). Whole mount *in situ* hybridization for *bax* during the first 60 hours post amputation confirm the expression dynamics and indicates that similar to other genes from the expression cluster R6 (Fig. 2 Bii) it is activated at the amputation site as early as 2hpa and progressively increases its expression in the mesenteries in later stages (Fig. 4Aii).

**Figure 4:**
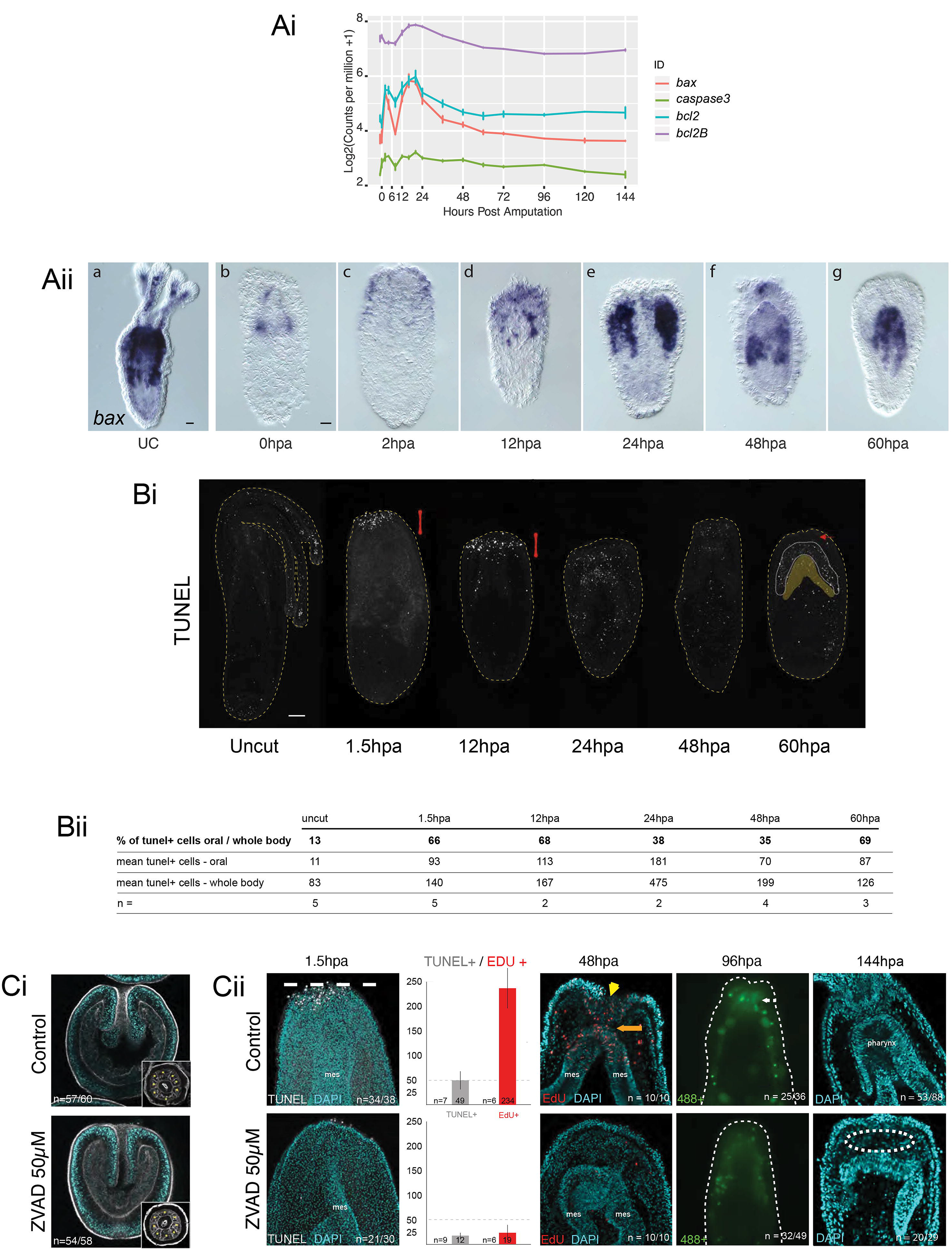
Apoptosis is required for regeneration, not embryogenesis. **(Ai)** Apoptosis genes (*bax, caspase3, bcl2,* and *bcl2B*) found in the regeneration specific module R-6 are activated transcriptionally early in response to injury. **(Aii)** Spatiotemporal expression pattern of *bax* during the first 60 hours post amputation by whole mount *in situ* hybridization. **(Bi)** TUNEL staining (white dots) in uncut controls and during regeneration (1.5hpa – 60hpa). The yellow dashed line indicates the outline of the polyp with the oral part / amputation site to the top. The red bar highlights the amputation site with increased TUNEL+ cells and the yellow patch indicates the tissues that link the mesenteries to the endodermal layer of the body wall (patch define by the white discontinued line). The red arrow designates the amputation site epithelia devoid of TUNEL+ cells. **(Bii)** Cell count and ratio of TUNEL positive cells (TUNEL+) detected at the amputation site *vs* the entire body. *n* indicates the number of analyzed animals. **(Ci)** Treatment with the pan-caspase inhibitor ZVAD does not affect embryonic development (48hpf). Embryos are stained with Phalloidin (f-actin in white) and DAPI (nuclei in turquoise). **(Cii)** Conversely zVAD treatment blocks regeneration at an early stage. In zVAD treated amputated polyp: TUNEL staining (1.5hpa: TUNEL, white) is inhibited, cell proliferation (48hpa: Edu, red) is also strongly reduced and auto-fluorescence of the pharynx does not re-emerge (96hpa: 488, green) indicating a failure of regeneration in those conditions. The morphology of zVAD treated amputated polyp at 144hpa (144hpa: DAPI, blue) shows lack of pharynx and tentacles, as well as the absence of the contact between the mesenteries and the epithelia of the amputation site indicating an early inhibition of the regeneration event.

We next performed a time series of TUNEL staining to examine the dynamics of apoptosis during embryogenesis and regeneration. While only few apoptotic cells can be observed during embryonic development (data not shown), during regeneration we observed a burst of apoptotic activity after amputation at the cut site as early as 1.5 hpa which perdured through 12hpa (Fig. 4Bi, Bii, Fig. S3). At 24hpa apoptotic activity is not detectable anymore at the wound site but randomly detected throughout the body, and at 60hpa, increasingly restricted around the mesenteries. Interestingly, this dynamic TUNEL profile (Fig. 4Bi, Bii) is reminiscent of the *bax* expression pattern (Fig. 4Aii).

To test whether or not apoptosis is indeed a regeneration specific process we used the pan-caspase inhibitor Z-VAD ^46^ to block apoptosis during embryogenesis and regeneration (Fig 4C, Fig. S4). *Nematostella* embryos treated continuously with Z-VAD after fertilization developed normally, showing no developmental defect (Fig. 4Ci) and metamorphosed on time (not shown). In contrast, regenerating *Nematostella* treated continuously with Z-VAD immediately after amputation were blocked in a very early regenerative stage (stage 0.5), preventing the physical interaction between the fused oral tip of the mesenteries and the epithelia of the wound site (Fig. 4Cii). Furthermore, amputated animals treated with ZVAD exhibited little to no cell proliferation, and never regenerated the pharynx or tentacles. This suggests an instructive function of apoptosis necessary for the induction of cell proliferation and the ensuring of a normal regenerative program (Fig. 4Cii). Thus, the present large-scale intra-species transcriptomic comparison of embryonic and regeneration datasets has enabled us to determine regeneration specific genes and processes, strongly suggesting that apoptosis in *Nematostella* is a regeneration-specific process.

### MEK/ERK signaling is required for the onset of regeneration

Our bioinformatics approach strongly suggests an important rearrangement of the genetic embryonic program during regeneration. Thus, we sought to functionally assess this observation by dissecting the MEK/ERK signaling pathway during the onset of regeneration and comparing the regeneration GRN with the GRN underlying embryonic development that is partly driven by MEK/ERK signaling ^22,47,48^. Using a monoclonal antibody directed against the phosphorylated/activated form of ERK (pERK, Fig S5A) in uncut animals and during the time course of regeneration, we detected localized pERK staining as early as 1hpa in the body wall epithelia of the amputation site as well as the oral tips of the mesenteries (Fig. 5A). After wound-healing (6hpa), this staining remains localized as regeneration progresses to reach a peak at 48hpa (Fig. 5A). A localized signal is detected in the reforming pharynx (72, 96hpa), after which the signal begins to be re-detected ubiquitously throughout the body (120, 144hpa), reminiscent of the staining detected in uncut controls (Fig. 5A).

**Figure 5:**
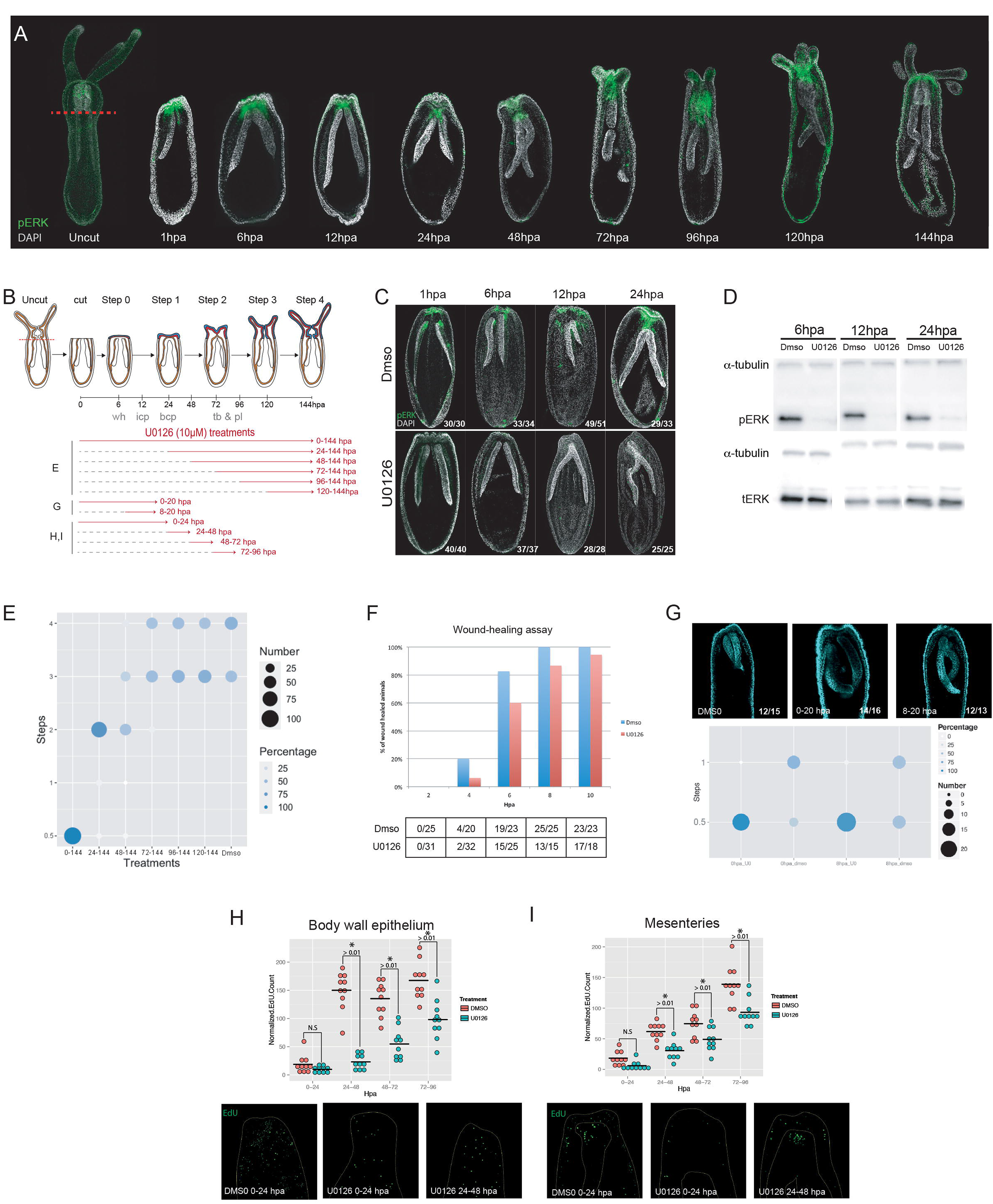
Inhibition of ERK phosphorylation by UO126 is required for regeneration. (A) Immunohistochemistry using and anti-pERK antibody revealing the activated/phosphorylated form of ERK (green) in uncut and regenerating polyps from 1 hpa to 144 hpa (counterstaining: DAPI). (B) Schematic representation of the various U0126 treatments that were applied during different windows of time during regeneration in the subsequent experiments. Polyps were fixed at the end of each treatments and analyzed. (C) Immunohistochemistry and Western blot (D) using and anti-pERK antibody (green) at various moments during regeneration in the absence or presence of the MEK inhibitor U0126 (Control condition: DMSO). In addition to anti-pERK, anti-tERK (total ERK) and anti-alpha-tubulin antibodies were used as controls for the western blot (D). (E) Phenotypic analysis resulting from UO126 treatments. The plot represents the regeneration step (x-axis) the treated polyps are in and the y-axis indicates the duration of U0126 treatments. 0.5 indicates the step following wound healing. The size of the dots and the blue shades represents the number of individuals per step and the percentage of polyps per treatment per step, respectively. (F) Diagram of results obtained from the compression/wound healing assays comparing UO126 treated polyps (red bars) during the first 10 hours of regeneration to DMSO controls (blue bars). (G) Confocal stacks of regenerating polyps counterstained with DAPI (cyan) focused at the mesenteries at 20 hpa after a 0-20 or 8-20 hpa treatment with UO126 and the dotplot shows the distribution of polyps between steps 0.5 and 1 according to the treatments. (H-I) Number of EdU+ cells (green) per polyps in the body wall epithelium (H) and the mesenteries (I), respectively. The x-axis shows the various 24 hours windows of treatment with UO126 (blue dots) or DMSO (red dots) and the y-axis show the number of EdU+ cells per polyp, represented by single dots (student t-test p.value <0.01).

Continuous treatment of bisected juveniles with the potent MEK inhibitor, UO126 ^49^, prevented activation of pERK in response to injury at the amputation site (Fig. 5B, C, D, S5B) and blocked regeneration at a very early step (step 0.5) preventing the reformation of the pharynx and tentacles when compared to 144hpa controls (Fig. 5E). When treatments were started at 24 or 48hpa, regeneration was blocked at step 2, and steps 2 and 3, respectively (Fig.5E). When U0126 was added at 72hpa or later time points, regeneration was completed successfully (steps 3 and 4) as in controls (Fig. 5E). These results indicate that MEK/ERK signalling is crucial for initiating a regenerative response (steps 0.5-1) as well as initiating the reformation of lost body parts (steps 2-3).

MEK/ERK signalling has been proposed to be required for wound-healing in *Nematostella* ^33^. Using an *in vivo* wound-healing assay ^31^, we determined that inhibition of MEK/ERK does not block, but delays wound-healing for 2-4 hours (Fig. 5F). To determine whether this delayed wound-healing is responsible for the arrest in the regeneration process, we compared the effects of U0126 treatments starting at 0hpa or 8hpa (*i.e*. after wound-healing is completed ^31^) on their capacity to reach step 1. While the majority of control animals (15/24) reached step 1, U0126 treated animals were consistently blocked at step 0.5 (Fig. 5G), indicating that regeneration is blocked independently of the delayed wound-healing process.

Cell proliferation in *Nematostella* is activated upon amputation and it is required for oral regeneration ^29,31^. To investigate whether MEK/ERK signalling is needed for the initiation or maintenance of cell proliferation, we treated bisected juveniles at various time frames (0-24hpa, 24-48hpa, 48-72hpa and 72-96hpa) during regeneration and quantified the number of cells in S-Phase (EdU+ cells) in the body wall epithelia (Fig. 5H) and the epithelia of the mesenteries (Fig. 5I) at the end of treatment. Surprisingly, following the same treatment setup as above (Fig, 5G, 0-24hpa, 8-24hpa) we did not observe any significant variation in cell proliferation between control and treatment in neither the localization, nor the amount of EdU+ cells (Fig. 5H,I, S6B). When treatments were applied at later time points (24-48hpa, 48-72hpa, 72-96hpa), the number of proliferating cells was significantly reduced in all conditions, when compared to the controls (Fig. 5H,I). Overall, these data indicate that MEK/ERK is not required to initiate, but rather to sustain cell proliferation during the regeneration process. These observations further indicate that the premature (step 0.5) arrest of regeneration upon inhibition of MEK/ERK is not related to cell proliferation but potentially through the inhibition of morphogenetic movements during the proliferation-independent phase of regeneration ^29,31^.

### Identification of synexpression groups at the onset of morphogenetic movements

We next decided to characterize the MEK/ERK-driven GRN underlying whole-body regeneration in *Nematostella* at 20hpa. This time point was chosen as it corresponds to the onset of morphogenetic movements (invagination/tissue reorganization, ^31^) as well as the likely onset of the regenerative program (Fig. 1C). Thus, this time point will allow a comparison with the previously defined embryonic GRN at the onset of the morphogenetic movements of gastrulation ^22,47,48^.

To identify potential MEK/ERK downstream targets responsible for launching the regenerative response we performed a differential gene expression analysis between 0 and 20hpa and identified 2,263 transcripts that were differentially expressed with an at least 2-fold variation (Table S4). We cross-referenced this list of transcripts with the genes that are part of the GRN underlying embryonic development ^22,47,48^ as well as the “regeneration-specific” genes identified above (Fig 1). This enabled us to identify 40 genes that are part of the embryonic GRN as well as 40 “regeneration-specific” genes that were upregulated at the onset of regeneration (Tables S5, S6, S7). From the 40 embryonic GRN genes, 25 have been described as downstream targets of MEK/ERK signalling ^22,47,48^, while the expression of 15 are controlled by the canonical Wnt (cWnt) pathway ^22,47,48^ at the onset of gastrulation.

We then performed an *in-situ* hybridization screen of 26 of these genes that we regrouped based on their spatial expression profile (Fig. 6Ai). Genes at 20hpa were either expressed in the Amputation site Ectodermis (AE: *wnt2, foxB, wntA* Fig. 6Aii a-c), Amputation site Gastrodermis (AG: *wnt7b, nkd1-like, wnt4, fz10*, Fig. 6Aii d-g), Amputation site Gastrodermis + Amputation site Mesenterial Tips (AG/AMT: *runt, moxD, axin-like, twist, wls,* Fig.6 Aii h-l; *erg*, *tcf,* Fig. 6Cdii,hii), Amputation site Gastrodermis + Mesenteries (AG/M: *sprouty, musk-like, porcupine-like, smad4-like*, Fig. 6Aii m-p; *carm, cad, egln1, bax,* Fig. 6Ciii-lii), Mesenteries (M, *hd50, mae-like, bicaudal-like, k50-5, pdvegfr-like, fox1, foxD1, mox1, phtf1-like,* Fig. 6Aii q-y) and the physa (P, *fgfA2*, Fig. 6Aii z). Taken together with their temporal expression profiles (Table S5, S6, S7), most of these genes form synexpression groups, potentially involved in the same biological process and underlying gene regulatory networks within their respective expression domains.

**Figure 6:**
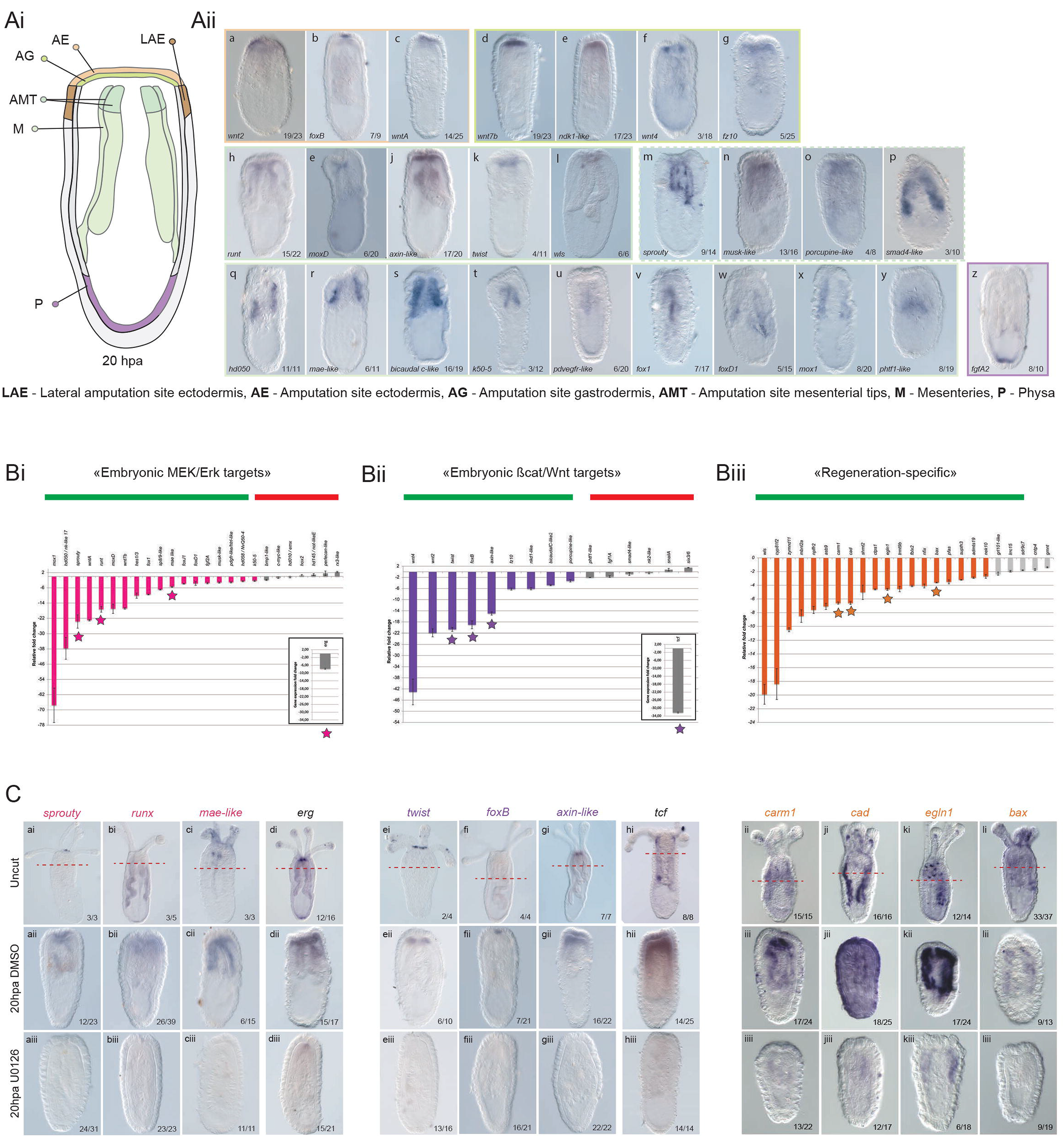
Identification and characterization of MEK/ERK downstream targets during regeneration. (Ai) Schematic representation of the various expression domains (*LAE – lateral amputation site ectodermis, AE - amputation site ectodermis, AG – amputation site gastrodermis, AMT – amputation site mesenterial tips, M - Mesenteries* and *P - physa*) characterizing the regenerating polyp as revealed by whole-mount *in situ* hybridization at 20 hpa. (Aii) *in situ* hybridization expression patterns of indicated genes transcriptionally up regulated at 20hpa. The colored rectangles surrounding the brightfield images, correspond to the expression pattern color code indicated in (Ai). The number in each panel indicates the total number of polyps with the represented pattern. (Bi-Biii) RT-qPCR analysis at 20 hpa comparing DMSO and UO126 treated polyps with genes identified as being transcriptionally upregulated at 20hpa. The analyzed genes corresponds either to (Bi) “embryonic MEK/ERK targets”, Bii) embryonic cWNT targets”, or (Biii) to regeneration-specific genes (Biii). The bold green line in each panel indicates genes that were at least 2-fold downregulated by U0126 treatments, while the bold red line indicates genes unaffected by the inhibition of MEK/ERK signaling during regeneration. The inserts indicate the RT-qPCR for *erg* (Bi) and *tcf* (Bii), the potential effectors of the MEK/ERK and cWNT pathways respectively. Genes with a star were further analyzed by whole mount *in situ* hybridization following U0126 treatments (C). The 1^st^ row corresponds to uncut DMSO treated controls with the amputation site indicated in dashed orange lines. The 2^nd^ row corresponds to DMSO treated controls at 20hpa and the 3^rd^ row are UO126 treated regenerating polyps fixed at the same time (20hpa). Numbers indicate the ratio of polyps with the shown expression pattern / total polyps.

### Regeneration activates a rewired embryonic gene regulatory network

To assess whether the above-identified genes are downstream targets of MEK/ERK during regeneration, we treated animals continuously after sub-pharyngeal amputation or starting at 8hpa (after wound-healing, Fig. S7) with U0126 and analysed their gene expression levels by RT-qPCR at 20hpa (Fig. 6B, S7). Strikingly, 19 out of the 25 identified embryonic MEK/ERK downstream targets (Table S5) are downregulated by U0126 treatments during regeneration (Fig. 6Bi). Similarly, 11 out of the 15 identified embryonic cWnt downstream targets (Table S6) are also downregulated following U0126 treatment (Fig. 6Bii). Finally, for the “regeneration-specific” pool of genes we only assessed 26 out of the 40 identified genes (Table S7) of which 19 were downregulated when MEK/ERK signalling was blocked by U0126 (Fig. 6Biii). We also assessed the effects of U0126 on *erg* (inset Fig. 6Bi) and *tcf* (inset Fig. 6Bii), two transcription factors that are effectors of MEK/ERK and cWnt, respectively, during embryonic development in *Nematostella* ^22,47^. Interestingly, we observed a reduced expression for *erg*, suggesting a negative feedback loop, and a strong down-regulation of *tcf* under those conditions. *In situ* hybridization of uncut, 20hpa DMSO treated and U0126 treated juveniles for a subset of these genes (*sprouty, runt, mae-like, erg*; *twist, foxB, axin-like, tcf*; *carm1, cad, egln1, bax)*, confirmed the RT-qPCR data (Fig. 6C). This analysis also revealed that at the onset of regeneration, MEK/ERK signalling controls expression of genes in several domains of the amputation site, including the AE, AG, AMT and M.

Taken together these data enabled us to propose a first MEK/ERK-dependent GRN blueprint underlying whole-body regeneration in *Nematostella* (Fig. 7A). This GRN indicates the potential direct or indirect downstream targets of MEK/ERK signalling at the onset of regeneration, that i) belong either to the group of embryonic cWNT or embryonic MEK/ERK targets and ii) are interconnected with “regeneration-specific” genes. During embryonic development, at the onset of gastrulation, cWNT activates first a set of downstream targets that is distinct from the ones controlled in a second phase by MEK/ERK signalling ^47^ (Fig. 7B). During regeneration however, MEK/ERK signalling appears to be activated shortly upon amputation potentially activating the cWNT pathway in a second phase (Fig. 7B). This functional approach thus confirmed the observation resulting from our cluster comparison (Fig. 3) suggesting that the GRN underlying regeneration follows a regeneration-specific network logic. This newly established network logic integrates distinct embryonic network modules (MEK/ERK and cWNT) and regeneration-specific elements to rapidly reform lost body parts (Fig. 7B).

**Figure 7:**
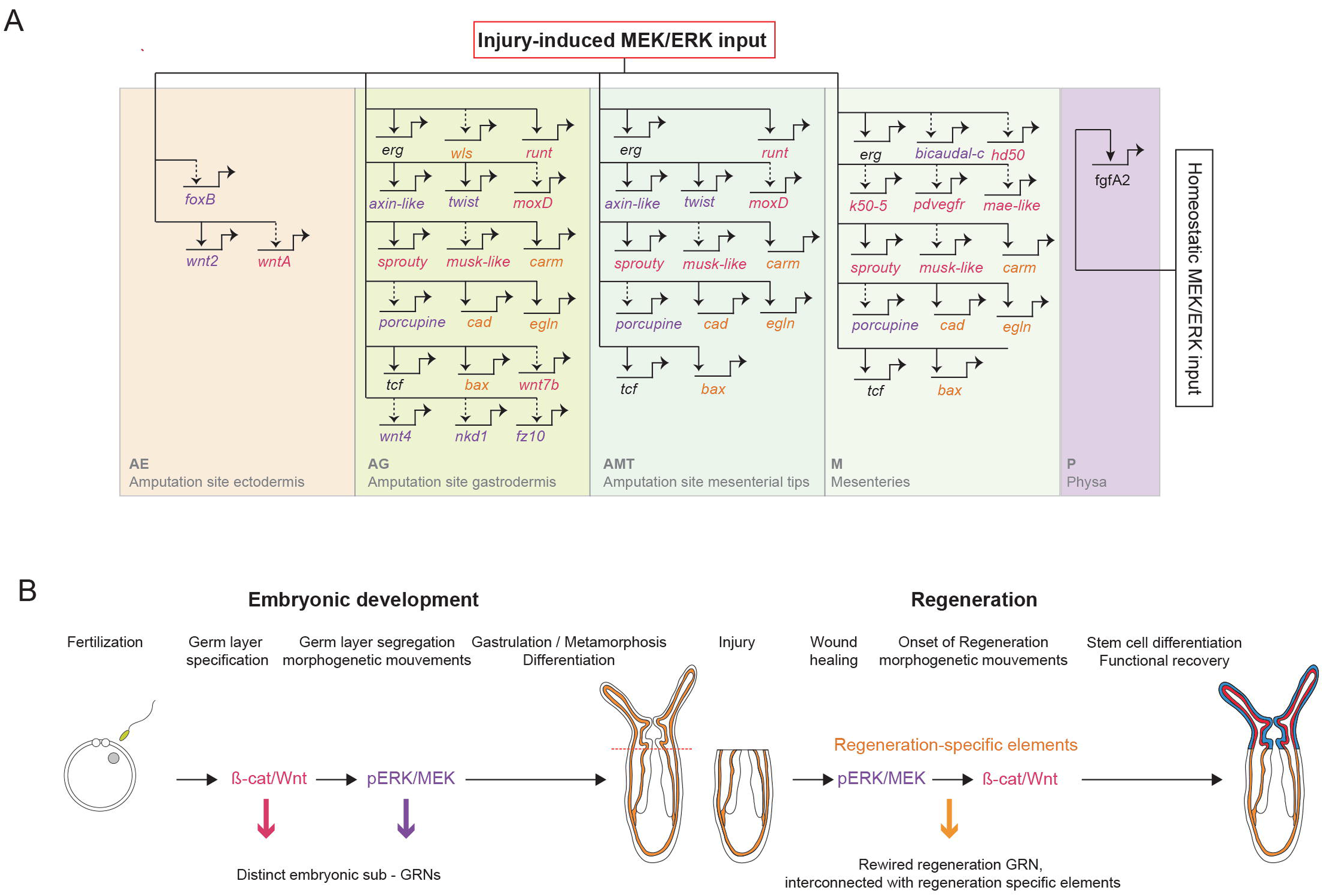
Blueprint of the MEK/ERK-dependent GRN underlying the onset of regeneration. (A) schematic representation of the MEK/ERK dependent gene regulatory network module at the onset of regeneration (20hpa). Colors of the boxes represent the expression patterns as shown in Fig. 6A. Genes are placed within this GRN according to their expression patterns in either one or several boxes. Solid lines indicate functional evidence that MEK/ERK signaling controls expression of the downstream target obtained by RT-qPCR and verified by *in situ* hybridization. No assumption on whether theses interactions are direct or indirect is made. (B) Global summary representing the results of the current study comparing the GRN underlying embryonic development and regeneration, highlighting that a rewired embryonic gene regulatory network interconnected with regeneration-specific elements is required to reform missing body parts following oral amputation in *Nematostella*.

## Discussion

### Regeneration is transcriptionally modest compared to embryonic development

In this work we used whole genome transcriptomic profiling to identify shared embryonic and regeneration-specific gene signatures. Comparing dynamically expressed genes in both processes, this approach revealed that regeneration is transcriptionally modest compared to embryogenesis (Fig. 1). This might be considered in contrast to a recent study carried out in sponges that showed that a similar number of genes is dynamically expressed during regeneration and post-larval development ^16^ However, it is important to point out that our study included early developmental stages and not only those following larval development, therefore covering a larger set of developmental processes. This might be the reason underlying the drastic difference of transcriptional dynamics between embryonic development and regeneration in *Nematostella*. We therefore favour the interpretation that regeneration only partially re-activates the embryonic program in response to the amputation stress, reflecting the largely differentiated cellular environment of the injury site. Interestingly, this is also confirmed by the findings that genes associated with the embryonic MEK/ERK (25/88) and cWNT (15/33) GRN modules are only partially re-deployed during regeneration (Table S3, S4) ^22,47^.

### Apoptosis is a and regeneration-specific process

This global comparison of genes dynamically expressed during embryonic development and regeneration revealed a set of genes whose expression dynamics are specific to regeneration and includes genes associated with apoptosis (Fig. 1). Similarly, by comparing embryonic and regenerative gene expression modules, we identified a gene module deployed early in regeneration that involves apoptosis (Fig. 3). These genes belong to so-called pro-(e.g. *bax, caspase-3*) and anti- (e.g. *bcl2, bcl2b*) apoptotic factors, highlighting the importance of a fine-balanced regulation of programed cell death during regeneration. However, the origin of the cells and whether those factors are activated in the same or neighboring cells remains to be elucidated.

Apoptosis has initially been associated with late developmental processes such as digit formation in mammalians ^50^ and tail regression during metamorphosis in ascidians ^51^. Following injury, apoptosis plays also an important role in various metazoans including the freshwater polyp *Hydra* ^52^, *Drosophila* ^53^ and Zebrafish ^54^. Using Z-VAD, a pharmacological pan-caspase inhibitor, classically used to block apoptosis in a variety of research models ^55^, we show that apoptosis is a cellular process specific to whole body regeneration in *Nematostella,* when compared to embryonic development (Fig. 4). When apoptosis is blocked, tissue-contact between the mesenteries and the amputation site as well as proliferation are prevented. Thus, we conclude that not only the classically described “destructive” function, but also the “instructive” function of apoptosis ^56^ are required to trigger a fine-tuned regenerative response in *Nematostella.* While apoptosis might be important for regeneration in general, the regeneration-specific role of apoptosis we highlight in this study, may represent a process common to all whole-body regenerators. Thus, regeneration is a partial re-use of the embryonic genetic programs but with important differences in its activation, which in the case of *Nematostella*, depends on apoptotic signals.

### Distinct implications of MEK/ERK signalling during regeneration

In order to experimentally verify the outcome of our global comparative cluster analysis and the prediction that the regeneration GRN follows a reshuffled network logic, we focussed on the MEK/ERK signalling pathway. Like a variety of other metazoans including *Hydra*, planarians, zebrafish and other and vertebrates ^57–61^ this signalling pathway seems not required for wound-healing but is crucial for regeneration in *Nematostella* (Fig. 5). In fact, we found that MEK/ERK signalling is essential for tissue remodelling, initiating the regeneration GRN and maintaining (not activating) proliferation. Importantly, these results are in line with our previous observations describing proliferation-independent (wound-healing and initiation of regeneration up to stage 1) and proliferation-dependent phases (reformation or lost body parts) of the wound-healing/regeneration process in *Nematostella* ^31^. Further analysis of the upstream activator of MEK/ERK and the downstream effectors in each context will contribute to have a better understanding of how this single pathway is reused in multiple regenerative contexts.

MEK/ERK signalling has been shown to activate programmed cell death (apoptosis) in *Hydra*, causing the release of Wnt3 and the induction of cellular proliferation at the amputation site ^52,62^. In a similar manner in *Nematostell*a, puncture ^33^ or sub-pharyngeal amputation (this study) induces apoptosis shortly after injury. However, inhibiting MEK/ERK signalling in *Nematostella* in either of the two wounding conditions has no visible effects on the injury-induced activation of apoptosis ^33^ (data not shown). These observations highlight that MEK/ERK signalling in *Nematostella* acts independently of apoptosis. This is in line with our results showing that apoptosis is necessary for the onset of proliferation, while MEK/ERK is not required for the onset but the maintenance of proliferation during regeneration (Fig. 5). While this observation is different from *Hydra* ^52,62^, it is strikingly similar to what has been observed in planarians ^57–61^ suggesting a conserved apoptosis-independent role of MEK/ERK to initiate and maintain the regeneration program following wound-healing. Further studies using additional whole body regeneration models are required to gain additional insight into the evolution of the role(s) of MEK/ERK signalling in launching the reformation of lost body parts.

### Regeneration deploys a novel network logic to activate the regenerative process

In the present study we used a combination of an unbiased large-scale bioinformatics approach and a signalling pathway targeted approach to compare embryonic development and regeneration at the gene regulatory level. Doing so, we suggest and experimentally confirm that the regeneration GRN is a partial and reshuffled redeployment of embryonic GRN modules interconnected with regeneration-specific elements (Fig. 7). In fact, during embryonic development cWNT and MEK/ERK activate a distinct set of downstream targets ^22,47,48^, while during regeneration MEK/ERK signalling is able to activate not only embryonic MEK/ERK targets but also embryonic cWNT as well as “regeneration-specific” genes. At this stage, we cannot provide details on the nature of the regulatory inputs (direct or indirect) of the effector of MEK/ERK on these downstream targets, thus further studies looking at chromatin accessibility during regeneration are required. This is particularly important to identify regeneration-specific enhancers ^63,64^ that may stimulate a regenerative response in tissues that have lost this capacity. A recent study assessing the latter during regeneration in the acoel *Hofstenia miamia* has revealed that the transcription factor Egr (Early growth response) is activated early upon injury and controls the expression of a large set of downstream targets ^65^. While Egr may play a similar role in planarians ^66^ and potentially in sea stars ^66^, its expression pattern in *Nematostella* (NvERTx.4.69506, nvertx.ircan.org) does not support an evolutionarily conserved regeneration-induction function in cnidarians. However, another transcription factor, runx/runt ^57,65,66^, this study) might be a conserved key player at the onset of regeneration within metazoans (ex. Sea star, acoel, *Hydra, Nematostella*), whose expression in response to injury is regulated by MEK/ERK signalling.

The combination of comparative transcriptional profiling and signaling pathway based functional assays shown in this study, clearly highlights the utility in considering not just individual gene use but how those genes are arranged into co-expression and gene regulatory modules. The identification of expression clusters along with a set of “regeneration-specific” genes provide valuable information to identify, in the future, regeneration-specific enhancers that drive injury-induced expression. Further studies, especially those comparing the activation of the regeneration GRNs across species, will provide novel insight into our understanding of why certain organisms can regenerate while others cannot. Furthermore, identification of key elements involved in re-deployment of signaling pathways can unlock hidden regenerative potential in poorly regenerating organisms.

## Acknowledgements

We thank Marina Shkreli (IRCAN), Eric Gilson (IRCAN) and Gianni Liti (IRCAN) for suggestions and critical reading of the manuscript as well as Valérie Carlin for animal husbandry and care. The authors also acknowledge the IRCAN’s Molecular and Cellular Core Imaging (PICMI) Facility. PICMI was supported financially by FEDER, Conseil régional Provence Alpes-Côte d’Azur, Conseil départemental 06, Cancéropôle PACA, Gis Ibisa and INSERM.

This work was supported by an ATIP-Avenir award (Institut National de la Santé et de la Recherche & Centre National de Recherche Scientifique) funded by the Plan Cancer (Institut National du Cancer, C13992AS), Seventh Framework Programme (CIG #631665), Fondation ARC pour la Recherche sur le Cancer (PJA2014120186), the French Government (National Research Agency, ANR) through the “Investments for the Future” programs LABEX SIGNALIFE ANR-11-LABX-0028 and IDEX UCAJedi ANR-15-IDEX-01 to **E.R**. as well as the Fondation ARC (PDF20141202150) to **J.F.W.**, Fondation pour la Recherche Médicale to **A.R.A** (SPF20130526781), **J.E.C** (SPF20170938703) and **H.J** (#FDT20170437124), the Ministère de l’enseignement Supérieur et de la Recherche to **H.J.** and la Ligue contre le Cancer to **K.N.**

## Materials and Methods

### Animal culture, spawning, embryo rearing, and amputation

Adult *Nematostella vectensis* were cultured at 16°C in the dark in 1/3 strength artificial sea water (ASW) as previously described (Amiel et al. 2017). The day before spawning the animals were fed with oysters and were then transferred to a light table for 12 hours. For embryology experiments, embryos were cultured at 18°C in the dark in 1/3 strength ASW until desired timepoint. Regeneration experiments were performed using six weeks-old juveniles raised at 22°C in the dark in 1/3 strength artificial sea water (ASW) as previously described (Amiel et al. 2017). Amputations were performed by first relaxing the juveniles in MgCl_2_ followed by sub-pharygneal amputation.

### RNA extraction, sequencing, read mapping and quantification

Detailed methods of the RNAseq methodology can be found in (Warner et al. 2018) and are described briefly here. RNA from ∼250 embryos was extracted as previously described at 24, 48, 72, 96, 120,144, 168, 172, and 196hpf in duplicate^34^. Additionally, RNA from ∼350 six weeks-old juveniles was extracted as previously described at uncut, 0, 2, 4, 8, 12, 16, 20, 24, 36, 48, 60, 72, 96, 120, and 144 hours post amputation in triplicate (Warner et al. 2018). cDNA libraries were prepared using the TruSeq stranded mRNA Library Prep Kit (Illumina #-NP-202-1001) and sequenced on an illumina NextSeq500 sequencer. Additional embryonic datasets were obtained from two previously published studies originally reported in Fischer et al. ^35^ (Illumina HiSeq 100bp paired end replicates sampled hourly from 0-19 hours post fertilization), and a second embryonic dataset originally reported in Helm et al. 2013 ^36^ (NCBI short read archive Project: PRJNA189768; Illumina HiSeq 50bp single end replicates sampled from 2, 7, 12, 24, 120 and 240). All reads from each dataset were processed equivalently. Reads were first quality filtered and adapter trimmed using trimmomatic ^67^ and cutadapt ^68^. Single end reads for each dataset, regeneration and the three embryonic datasets Fischer, Helm, and Warner, were aligned to a transcriptome assembly comprised of both embryonic and regeneration RNAseq data using Bowtie2 ^69^ and read counts were quantified using RSEM ^70^. Gene level counts were obtained by merging transcripts with the same top BLASTn hit to the Nemve1 filtered gene models ^34^. Gene models that did not have >5 counts in at least 25% of the samples, were excluded. Each dataset was then normalized separately using the R package edgeR and the counts per million (cpm) mapped reads were calculated ^71^. Batch effects of the embryonic datasets were corrected using the R function ComBat from the SVA package using timepoint as a categorical covariate (Fig S1) ^37^. For all downstream calculations only embryonic timepoints from 7hpf (the estimated onset of zygotic transcription) onward are considered so that only gene transcription dynamics are measured and compared.

### Identification of regeneration specific genes and GO-term enrichment

For each dataset we calculated differential expression for each Nemve1 gene model using edgeR and comparing each time point to t_0_ (t_0_= 7hpf Helm and Fischer dataset; 24hpf Warner dataset; and 0hpa for the regeneration dataset) ^34–36^. We define a significantly differentially expressed gene as having an absolute log_2_(Fold Change) > 2 and a false discovery rate (FDR) < 0.05. These differentially expressed gene lists were compared to identify overlapping and regeneration specific genes. GO term enrichment of the regeneration specific gene list was calculated using a Fisher’s exact test and the R package topGO on the GO terms identified from comparing the Nemve1 gene models to the UniProt databases Swissprot and Trembl using the BLASTx like program PLASTx (evalue cutoff 5e-5) ^72^. All identified GO terms were used as a background model. The resulting GO term list was reduced and plotted using a modified R script based on REVIGO ^73^.

### Embryonic versus Regeneration dataset comparison: Principal component analysis, Fuzzy c-means clustering and cluster conservation

The ensuing analyses were performed using log_2_(cpm+1) transformed gene-level quantification (Nemve1 filtered gene models). The expression profiles for each Nemve1 gene model were clustered using the R package mFuzz ^40^ on the combined embryonic dataset and the regeneration dataset separately. The cluster number was set to 9 for the regeneration data and 8 for the embryonic datasets as these numbers produced well-separated clusters with minimal overlap (Fig. S2) and represent the inflection point at which the centroid distance between clusters did not significantly decrease with the addition of new clusters (Fig. S2). Genes that did not have a membership score above 0.75 were considered noise and designated as cluster 0. GO-term enrichment testing for each cluster was performed as described above. Cluster overlap was calculated for genes that were detectable in both datasets using the function overlapTable from the R package WGCNA using the regeneration cluster assignments as the reference set. A zStatistic of cluster preservation was also calculated using the function coClustering.permutationTest from the WGCNA package using the regeneration cluster assignments as the reference set and 1000 permutations.

### In situ hybridization

Whole-mount *in situ* hybridization was performed as previously described ^74^. The probes used in this study were synthetized and labeled with digoxigenin according to the protocol described in ^47^, and were diluted to 0.1 ng/µl in fresh hybridization solution. Anti-Dig/AP used at 1:5000 in blocking solution and incubated at 4°C overnight.

### Apoptotic cell death staining

After relaxing *Nematostella* polyps in MgCl_2_ for 10-15 minutes, animals were fixed in 4% paraformaldehyde (Electron Microscopy Sciences # 15714) in 1/3 ASW during 1 hour at 22°C or overnight at 4°C. Fixed animals were washed three times in PBT 0.5% (PBS1x + Triton 0.5%). To detect cell death the “In Situ Cell Death AP kit” (Roche, #11684809910) was used. The manufacturer protocol was modified as follow: 1) Fixed animals were permeabilized using 0.01mg/ml Proteinase K for 20min at 22°C; 2), Washed twice in PBS1x; 3) Refixed in 4% paraformaldehyde in PBS1x for 1 hour at 22°C; 4) Washed 5 times in PBS1x; 5) Incubated with 50μL of TUNEL reaction mixture for one hour (Roche protocol); 6) Washed 5 times in PBS1x; 7) Fixed animal were observed for 488 fluorescence. Positive (DNase I treatment after step 4) and negative (without TUNEL-Enzyme) controls were obtained using the manufacturer’s “In Situ Cell Death AP kit” protocol.

### Pharmaceutical drug treatments to block apoptotic cell death

Apoptotic cell death was blocked using the pan-caspase inhibitor Z-VAD-FMK (named ZVAD throughought the manuscript, #ALX-260-020-M001, Enzo Life Sciences Inc, Farmingdale, NY, USA). A stock solution at 10mM in DMSO was prepared for Z-VAD, kept at -20°C and diluted in 1/3 ASW at a final concentration of 10µM or 50µM prior to each experiment (Fig. S4). Each Z-VAD treatment was performed in a final volume of 500µl 1/3X ASW in a 24 well plate using the adequate controls (1/3 ASW or DMSO). Reagents were changed every 24h to maintain activity for the duration of the experiments.

### Pharmaceutical drug treatments to block apoptotic cell death and MEK inihibition

Apoptotic cell death was blocked using the pan-caspase inhibitor Z-VAD-FMK (named ZVAD through the manuscript, #ALX-260-020-M001, Enzo Life Sciences Inc, Farmingdale, NY, USA). A stock solution at 10mM in DMSO was prepared for Z-VAD, kept at -20°C and diluted in 1/3 ASW at a final concentration of 10µM or 50µM prior to each experiment (Fig. S4). Each Z-VAD treatment was performed in a final volume of 500µl 1/3 ASW in a 24 well plate using the adequate controls (1/3 ASW or 0.1% DMSO in 1/3 ASW). Reagents were changed every 24h to maintain activity for the duration of the experiments.

Phosphorylation/activation of ERK was prevented by the MEK inhibitor, U0126 (Sigma U120-1MG). A stock of 10 mM in DMSO was prepared and stored and - 20°C. Each treatment with U0126 were performed with a final concentration of 10 μM in 1/3 ASW. Treatments were carried the same way as ZVAD treatments.

### Imaging the effect of U0126 treatments during regeneration

The regenerating *Nematostella* polyps were fixed as described in *Apoptotic cell death staining*. For regeneration step characterization, we counterstained the nuclei and actin filament with Hoechst (Invitrogen #33342, Carlsbad, CA, USA) and BODIPY FL PhallAcidin 488 (Molecular Probes #B607, Eugene, OR, USA), respectively. Hoechst was diluted to 1/5000 and BODIPY FL PhallAcidin 488 was diluted to 1/200 in PBS. The phosphorylated form of ERK was visualized using a monoclonal antibody directed against its active form, p-ERK (Cat.#4377; Cell Signaling Technology) and the protocol used was described in ^33^. For cell proliferation analysis we used the Click-it EdU kit (Invitrogen #C10337 or #C10339, Carlsbad, CA, USA) following the protocol from ^29^.

### Protein extraction and western blotting

The protein extracts were obtained from 15 adults per replicate and each experiment was performed in triplicate. The animals were placed in 1.5 ml eppendorff tubes and spun on bench centrifuge to remove all 1/3 ASW before adding 300μl of lysis buffer (HEPES 50mM, NaCl 150mM, NaF 100mM, EDTA 10mM, NA4P207 10mM). Then animals were sonicated 5 x 10 sec with incubation on ice between each round. After sonication 200μl of Lysis buffer complemented with 1% Triton X-100 and a protease inhibitor cocktail (Apoprotine 20μg/ml, Vanadate 1mM, AEBSF 250μg/ml, Leupeptine 5μg/ml) was added. The samples were then centrifuged for 20 minutes at 12,000 rpm at 4°C before collecting the supernatant into new tubes. The BC assay protein quantification kit (Interchim Upima, 40840A) was used to quantify the protein concentration. Samples were subsequently aliquoted by mixing 75μl of protein extract with 25μl of 4X Laemmli buffer (Bio-Rad #1610747) and stored at -20°C. Electrophoresis was carried out in 7.5% SDS polyacrylamide gel and samples were denatured at 85°C for 5 min before loading. For each sample we used 30μg of protein and followed the protocol described in ^75^. Briefly, migration was carried in migration buffer (Tris 3g/L, Glycine 14.2g/L, SDS 1%) and transfer to the nitrocellulose membrane was performed in transfer buffer (Tris 3g/L, Glycine 14,4g/L, 20% Ethanol). Then the nitrocellulose membrane was saturated with Salin buffer (Tris 0,24g/L, NaCl 1,63g/L, 5%cBSA, 0.5% Tween-20) and incubated 1 hour at room temperature with 1/2500 of primary antibody anti-pERK (Cat.#4377; Cell Signaling Technology) diluted saline buffer. Revelation was carried out by chimioluminescence (EMD Millipore™ Substrats de chimioluminescence HRP Western Luminata™) on a chemioluminescence Imaging –Fusion SL (Vilber).

### RT-qPCR

RNA Extraction and RT-qPCR were performed following protocol described in ^22^. Briefly, we used 160 juveniles per replicate and each experiment was performed in three biological replicates. We used a 7900HT Fast Real-Time PCR System with 384-Well Block Module (Applied Biosystem) with Faststart universal SYBR Green Master (rox) (FSUSGMMRO Roche). The relative fold change for each gene expression was calculated by comparing treated and untreated conditions, where *gapdh* was used to normalized fold change (FC). The full list of qPCR primer pairs and their efficiency used in this study can be found in Table S8 (Regeneration specific genes) and in ^48^ (Embryonic genes).

## Supplementary figures

**FigureS1: Embryonic sample tree pre, post batch correction.**

Sample trees of embryonic log_2_(cpm+1) before batch correction (**top**) and after (**bottom**)

**Figure S2: Fuzzy c-means clustering minimum centroid distance and cluster overlap.**

(**Top**): Minimum centroid distances for embryogenesis (**left**) and regeneration (**right**) clusters. (**Bottom**): Overlap plots for embryogenesis (**left**) and regeneration (**right**) clusters show principal component analysis of the cluster centers. The overlap is visualised by lines with variable width indicating the strength of the overlap.

**Figure S3: TUNEL Assay controls.**

Close-up of the physa (a, a’) and the tentacles (b, b’) for the TUNEL assay positive control using DNAse I treatment. Uncut polyp (c, c’) and 1.50hpa (d, d’) for the TUNEL assay negative control without Enzyme. Green dashed arrows (b’, c’) indicate non-specific auto-fluorescent staining. n=[number of specimen with represented phenotype]/[total number of analyzed specimen]

**Figure S4: ZVAD dose-response assay and effects on tissue organization.**

(**A)** zVAD dose-response assay on juvenile (a, b, c) *vs* adult (a’, b’, c’) amputated polyps. Untreated polyps at 5 days post amputation (5dpa) (a, a’), 10µm zVAD treated polyp at 5dpa (b, b’) and 50µm zVAD treated polyp at 5dpa (c, c’). 50µm zVAD treatment induces the inhibition of regeneration in nearly 100% of the total number of cases (75 out of 77 polyps, c), while 10µm zVAD treatment fully inhibits regeneration in only half of the total number of cases (31 out of 61 polyps, b). **(B)** zVAD treatment does not affect tissues organization. Untreated tissues (a, a’, b, b’) and 50µm zVAD treated tissues (c, c’, d, d’) at 5dpa. Nucleus DNA staining (DAPI) is in blue (a, b, c, d). Cell membrane and muscle network actin filaments staining (phalloidin) are in white (a’, b’, c’, d’).

**Figure S5: WB DMSO + U0126 at various time points.**

Antibodies used are the same as in Fig. 5

**Figure S6: Effect of U0126 treatments at different time points.**

**Figure S7: RT-qPCR results comparison long (0-20hpa) vs short (8-20hpa) U0126 treatments, indicating that both treatments have the same effects on downstream targets.**

**Supplementary table 1: Regeneration specific genes**

**Supplementary table 2: GO-term enrichment of embryonic clusters**

**Supplementary table 3: GO-term enrichment of regeneration clusters**

## References

1. Bely, A. E. & Nyberg, K. G. Evolution of animal regeneration: re-emergence of a field. Trends in ecology & evolution 25, 161–170 (2010).

2. Morgan, T. H. Regeneration. Columbia University Biological Series. vol. 279 (1901).

3. Bryant, D. M. et al. A Tissue-Mapped Axolotl De Novo Transcriptome Enables Identification of Limb Regeneration Factors. CellReports 18, 762–776 (2017).

4. Habermann, B. et al. An Ambystoma mexicanum EST sequencing project: analysis of 17,352 expressed sequence tags from embryonic and regenerating blastema cDNA libraries. Genome Biology 5, R67 (2004).

5. Hutchins, E. D. et al. Transcriptomic analysis of tail regeneration in the lizard Anolis carolinensis reveals activation of conserved vertebrate developmental and repair mechanisms. PLoS ONE 9, e105004 (2014).

6. Rodius, S. et al. Analysis of the dynamic co-expression network of heart regeneration in the zebrafish. Scientific Reports 6, 26822 (2016).

7. Gardiner, D. M., Blumberg, B., Komine, Y. & Bryant, S. V. Regulation of HoxA expression in developing and regenerating axolotl limbs. Development (Cambridge, England) 121, 1731–1741 (1995).

8. Schaffer, A. A., Bazarsky, M., Levy, K., Chalifa-Caspi, V. & Gat, U. A transcriptional time-course analysis of oral vs. aboral whole-body regeneration in the Sea anemone Nematostella vectensis. BMC genomics 17, 718 (2016).

9. Millimaki, B. B., Sweet, E. M. & Riley, B. B. Sox2 is required for maintenance and regeneration, but not initial development, of hair cells in the zebrafish inner ear. Developmental Biology 338, 262–269 (2010).

10. Katz, M. G., Fargnoli, A. S., Kendle, A. P., Hajjar, R. J. & Bridges, C. R. The role of microRNAs in cardiac development and regenerative capacity. American journal of physiology. Heart and circulatory physiology 310, H528–41 (2016).

11. Imokawa, Y. & Yoshizato, K. Expression of Sonic hedgehog gene in regenerating newt limb blastemas recapitulates that in developing limb buds. Proceedings of the National Academy of Sciences 94, 9159–9164 (1997).

12. Carlson, M. R., Komine, Y., Bryant, S. V. & Gardiner, D. M. Expression of Hoxb13 and Hoxc10 in developing and regenerating Axolotl limbs and tails. Developmental Biology 229, 396–406 (2001).

13. Torok, M. A., Gardiner, D. M., Shubin, N. H. & Bryant, S. V. Expression of HoxD genes in developing and regenerating axolotl limbs. Developmental Biology 200, 225–233 (1998).

14. Özpolat, B. D. et al. Regeneration of the elbow joint in the developing chick embryo recapitulates development. Developmental Biology 1–10 (2012) doi:10.1016/j.ydbio.2012.09.020.

15. Wang, Y.-H. & Beck, C. W. Distal expression of sprouty (spry) genes during Xenopus laevis limb development and regeneration. Gene expression patterns : GEP 15, 61–66 (2014).

16. Soubigou, A., Ross, E. G., Touhami, Y., Chrismas, N. & Modepalli, V. Regeneration in sponge Sycon ciliatum partly mimics postlarval development. Development 147, dev.193714 (2020).

17. Czarkwiani, A., Dylus, D. V., Carballo, L. & Oliveri, P. FGF signalling plays similar roles in development and regeneration of the skeleton in the brittle star Amphiura filiformis. Development 148, (2021).

18. Sinigaglia, C. et al. Distinct gene expression dynamics in developing and regenerating limbs. Biorxiv 2021.06.14.448408 (2021) doi:10.1101/2021.06.14.448408.

19. Layden, M. J., Rentzsch, F. & Röttinger, E. The rise of the starlet sea anemone Nematostella vectensis as a model system to investigate development and regeneration: Overview of starlet sea anemone Nematostella vectensis. Wiley Interdiscip Rev Dev Biology 5, 408–428 (2016).

20. Hand, C. & Uhlinger, K. R. The Culture, Sexual and Asexual Reproduction, and Growth of the Sea Anemone Nematostella vectensis. The Biological Bulletin 182, 169–176 (1992).

21. Wikramanayake, A. H. et al. An ancient role for nuclear beta-catenin in the evolution of axial polarity and germ layer segregation. Nature 426, 446–450 (2003).

22. Röttinger, E., Dahlin, P. & Martindale, M. Q. A Framework for the Establishment of a Cnidarian Gene Regulatory Network for “Endomesoderm” Specification: The Inputs of ß-Catenin/TCF Signaling. Plos Genet 8, e1003164 (2012).

23. Leclère, L., Bause, M., Sinigaglia, C., Steger, J. & Rentzsch, F. Development of the aboral domain in Nematostella requires β-catenin and the opposing activities of six3/6 and frizzled5/8. Development (Cambridge, England) 143, 1766–1777 (2016).

24. Genikhovich, G. et al. Axis Patterning by BMPs: Cnidarian Network Reveals Evolutionary Constraints. CellReports 10, 1646–1654 (2015).

25. Wijesena, N., Simmons, D. & Martindale, M. Antagonistic BMP-cWNT signaling in the cnidarian Nematostella vectensis: Implications for the evolution of mesoderm. Mechanisms of Development 145, S114 (2017).

26. Reitzel, A., Burton, P., Krone, C. & Finnerty, J. Comparison of developmental trajectories in the starlet sea anemone Nematostella vectensis: embryogenesis, regeneration, and two forms of asexual fission. Invertebrate Biology 126, 99–112 (2007).

27. Burton, P. M. & Finnerty, J. R. Conserved and novel gene expression between regeneration and asexual fission in Nematostella vectensis. Development Genes and Evolution 219, 79–87 (2009).

28. Trevino, M., Stefanik, D. J., Rodriguez, R., Harmon, S. & Burton, P. M. Induction of canonical Wnt signaling by alsterpaullone is sufficient for oral tissue fate during regeneration and embryogenesis in Nematostella vectensis. Developmental dynamics : an official publication of the American Association of Anatomists 240, 2673–2679 (2011).

29. Passamaneck, Y. J. & Martindale, M. Q. Cell proliferation is necessary for the regeneration of oral structures in the anthozoan cnidarian Nematostella vectensis. BMC Developmental Biology 12, 1–1 (2012).

30. Bossert, P. E., Dunn, M. P. & Thomsen, G. H. A staging system for the regeneration of a polyp from the aboral physa of the anthozoan cnidarian Nematostella vectensis. Developmental dynamics : an official publication of the American Association of Anatomists 242, 1320–1331 (2013).

31. Amiel, A. et al. Characterization of Morphological and Cellular Events Underlying Oral Regeneration in the Sea Anemone, Nematostella vectensis. Int J Mol Sci 16, 28449–28471 (2015).

32. Amiel, A. R., Foucher, K., Ferreira, S., bioRxiv, E. R. & 2019. Synergic coordination of stem cells is required to induce a regenerative response in anthozoan cnidarians. biorxiv.org (2019) doi:10.1101/2019.12.31.891804.

33. DuBuc, T. Q., Traylor-Knowles, N. & Martindale, M. Q. Initiating a regenerative response; cellular and molecular features of wound healing in the cnidarian Nematostella vectensis. 12, 1–20 (2014).

34. Warner, J. F. et al. NvERTx: a gene expression database to compare embryogenesis and regeneration in the sea anemone Nematostella vectensis. Development (Cambridge, England) 145, dev162867 (2018).

35. Fischer, A. & Smith, J. Nematostella High-density RNAseq time-course. (2013) doi:10.1575/1912/5981.

36. Helm, R. R., Siebert, S., Tulin, S., Smith, J. & Dunn, C. W. Characterization of differential transcript abundance through time during Nematostella vectensis development. BMC genomics 14, 266 (2013).

37. Leek, J. T., Johnson, W. E., Parker, H. S., Jaffe, A. E. & Storey, J. D. The sva package for removing batch effects and other unwanted variation in high-throughput experiments. Bioinformatics 28, 882–883 (2012).

38. Adell, T., Saló, E., Boutros, M. & Bartscherer, K. Smed-Evi/Wntless is required for beta-catenin-dependent and -independent processes during planarian regeneration. Journal of embryology and experimental morphology 136, 905–910 (2009).

39. Bassat, E. et al. The extracellular matrix protein agrin promotes heart regeneration in mice. Nature 547, 179–184 (2017).

40. Kumar, L. & Futschik, M. E. Mfuzz: a software package for soft clustering of microarray data. Bioinformation 2, 5–7 (2007).

41. Fritzenwanker, J. H. J., Saina, M. M. & Technau, U. U. Analysis of forkhead and snail expression reveals epithelial-mesenchymal transitions during embryonic and larval development of Nematostella vectensis. Developmental Biology 275, 389–402 (2004).

42. Kusserow, A. et al. Unexpected complexity of the Wnt gene family in a sea anemone. Nature 433, 156–160 (2005).

43. Srivastava, M. et al. Early evolution of the LIM homeobox gene family. BMC biology 8, 4 (2010).

44. Niehrs, C. & Pollet, N. Synexpression groups in eukaryotes. Nature 402, 483–487 (1999).

45. Langfelder, P., Luo, R., Oldham, M. C. & Horvath, S. Is my network module preserved and reproducible? PLoS computational biology 7, e1001057 (2011).

46. Cikala, M., Wilm, B., Hobmayer, E., Böttger, A. & David, C. N. Identification of caspases and apoptosis in the simple metazoan Hydra. Curr Biol 9, 959–S2 (1999).

47. Amiel, A. R. et al. A bipolar role of the transcription factor ERG for cnidarian germ layer formation and apical domain patterning. Developmental Biology 430, 346–361 (2017).

48. Layden, M. J. et al. MAPK signaling is necessary for neurogenesis in Nematostella vectensis. BMC biology 14, 61 (2016).

49. DeSilva, D. R. et al. Inhibition of mitogen-activated protein kinase kinase blocks T cell proliferation but does not induce or prevent anergy. The Journal of Immunology 160, 4175–4181 (1998).

50. Suzanne, M. & Steller, H. Shaping organisms with apoptosis. Cell Death Differ 20, 669–675 (2013).

51. Chambon, J.-P. et al. Tail regression in Ciona intestinalis (Prochordate) involves a Caspase-dependent apoptosis event associated with ERK activation. Journal of embryology and experimental morphology 129, 3105–3114 (2002).

52. Chera, S. et al. Apoptotic Cells Provide an Unexpected Source of Wnt3 Signaling to Drive Hydra Head Regeneration. Developmental Cell 17, 279–289 (2009).

53. Milán, M., Campuzano, S. & García-Bellido, A. Developmental parameters of cell death in the wing disc of Drosophila. Proc National Acad Sci 94, 5691–5696 (1997).

54. Gauron, C. et al. Sustained production of ROS triggers compensatory proliferation and is required for regeneration to proceed. Scientific Reports 3, (2013).

55. Poss, K. D. Advances in understanding tissue regenerative capacity and mechanisms in animals. Nature Reviews Genetics 11, 710–722 (2010).

56. Ryoo, H. D. & Bergmann, A. The role of apoptosis-induced proliferation for regeneration and cancer. Cold Spring Harbor Perspectives in Biology 4, a008797–a008797 (2012).

57. Petersen, H. O. et al. A Comprehensive Transcriptomic and Proteomic Analysis of Hydra Head Regeneration. Molecular biology and evolution 32, 1928–1947 (2015).

58. Owlarn, S. et al. Generic wound signals initiate regeneration in missing-tissue contexts. Nature Communications 8, 2282 (2017).

59. Liu, P. & Zhong, T. P. MAPK/ERK signalling is required for zebrafish cardiac regeneration. Biotechnol Lett 39, 1069–1077 (2017).

60. Manuel, G. C., Reynoso, R., Gee, L., Salgado, L. M. & Bode, H. R. PI3K and ERK 1-2 regulate early stages during head regeneration in hydra. Development, Growth and Differentiation 48, 129–138 (2006).

61. Poss, K. D. et al. Roles for Fgf Signaling during Zebrafish Fin Regeneration. Dev Biol 222, 347–358 (2000).

62. Chera, S., Ghila, L., Wenger, Y. & Galliot, B. Injury-induced activation of the MAPK/CREB pathway triggers apoptosis-induced compensatory proliferation in hydra head regeneration. Development, Growth and Differentiation 53, 186–201 (2011).

63. Kang, J. et al. Modulation of tissue repair by regeneration enhancer elements. Nature 532, 201–206 (2016).

64. Goldman, J. A. & Poss, K. D. Gene regulatory programmes of tissue regeneration. Nature Reviews Genetics 11, 710–15 (2020).

65. Gehrke, A. R. et al. Acoel genome reveals the regulatory landscape of whole-body regeneration. Science (New York, N.Y.) 363, eaau6173–9 (2019).

66. Cary, G. A., Wolff, A., Zueva, O., Pattinato, J. & Hinman, V. F. Analysis of sea star larval regeneration reveals conserved processes of whole-body regeneration across the metazoa. BMC biology 17, 1–19 (2019).

67. Bolger, A. M., Lohse, M. & Usadel, B. Trimmomatic: a flexible trimmer for Illumina sequence data. Bioinformatics 30, 2114–2120 (2014).

68. Martin, M. Cutadapt removes adapter sequences from high-throughput sequencing reads. EMBnet Journal 17, 10–12 (2011).

69. Langmead, B., Trapnell, C., Pop, M. & Salzberg, S. L. Ultrafast and memory-efficient alignment of short DNA sequences to the human genome. Genome Biology 10, R25 (2009).

70. Li, B. & Dewey, C. N. RSEM: accurate transcript quantification from RNA-Seq data with or without a reference genome. BMC Bioinformatics 12, 323 (2011).

71. Robinson, M. D., McCarthy, D. J. & Smyth, G. K. edgeR: a Bioconductor package for differential expression analysis of digital gene expression data. Bioinformatics 26, 139–140 (2010).

72. Nguyen, H. V. & Lavenier, D. PLAST: parallel local alignment search tool for database comparison. BMC Bioinformatics 10, 329 (2009).

73. Supek, F., Bošnjak, M., Škunca, N. & Šmuc, T. REVIGO summarizes and visualizes long lists of gene ontology terms. PLoS ONE 6, e21800 (2011).

74. Genikhovich, G. & Technau, U. The starlet sea anemone Nematostella vectensis: an anthozoan model organism for studies in comparative genomics and functional evolutionary developmental biology. Cold Spring Harbor protocols 2009, pdb.emo129-pdb.emo129 (2009).

75. Gilmore, T., Wolenski, F. & Finnerty, J. Preparation of antiserum and detection of proteins by Western blotting using the starlet sea anemone, Nematostella vectensis. Protoc Exch (2012) doi:10.1038/protex.2012.057.

