## Supplementary figures and images for "Whole body regeneration deploys a rewired embryonic gene regulatory network logic"

### Suppl Figure 1

#### Pre Batch Correction

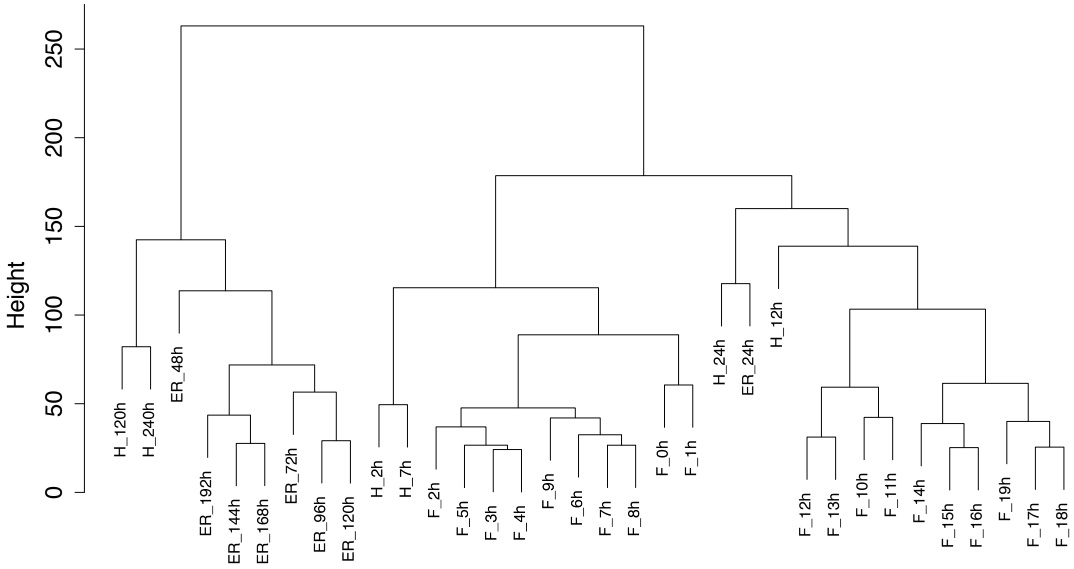


#### Post Batch Correction

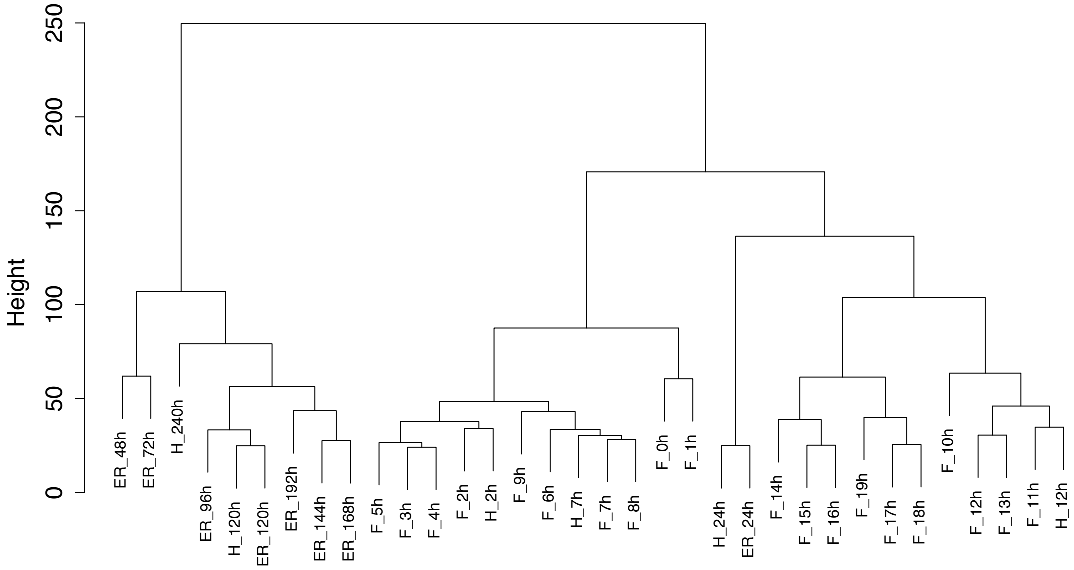

### Suppl Figure 2

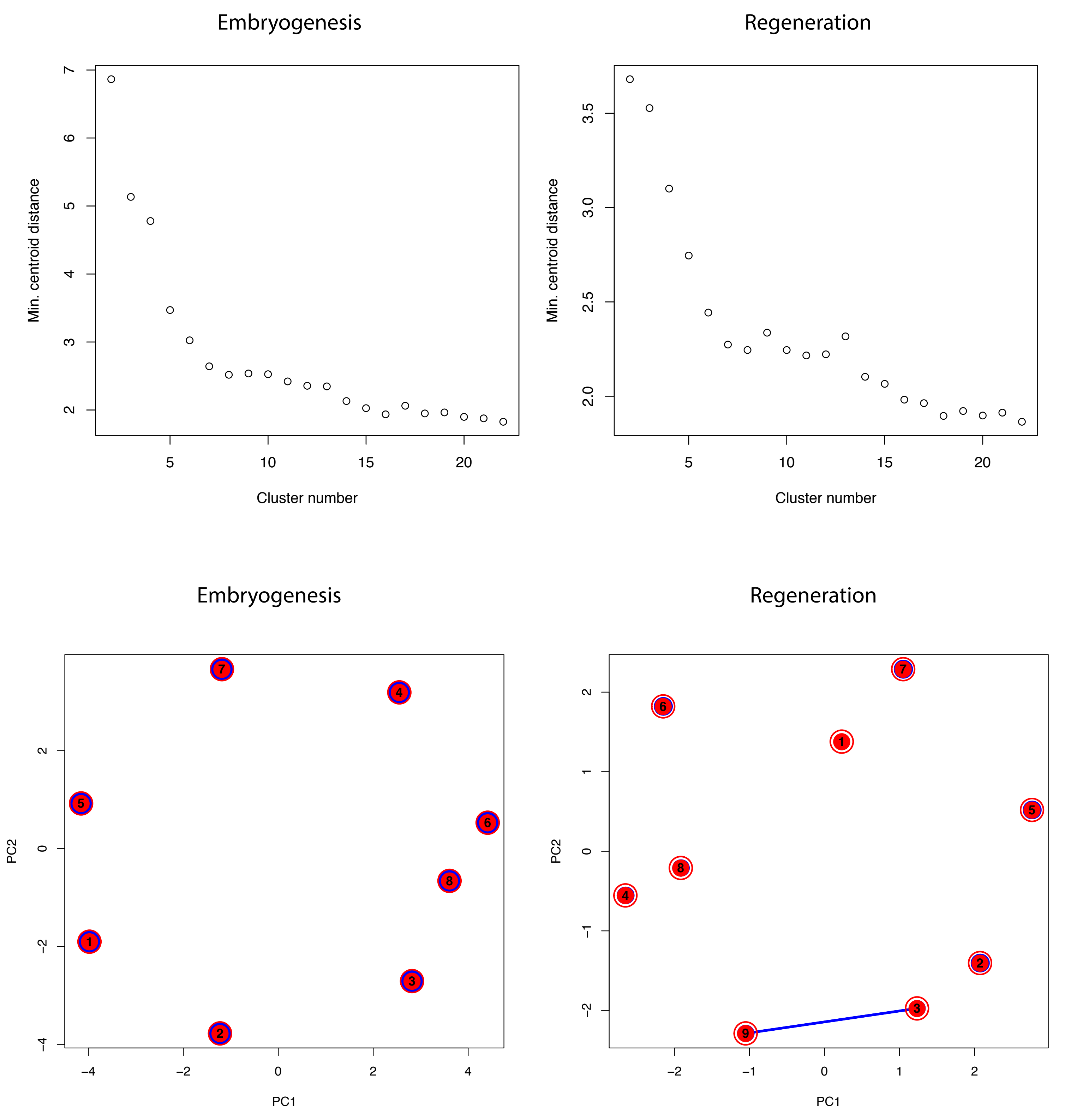

### Suppl Figure 3

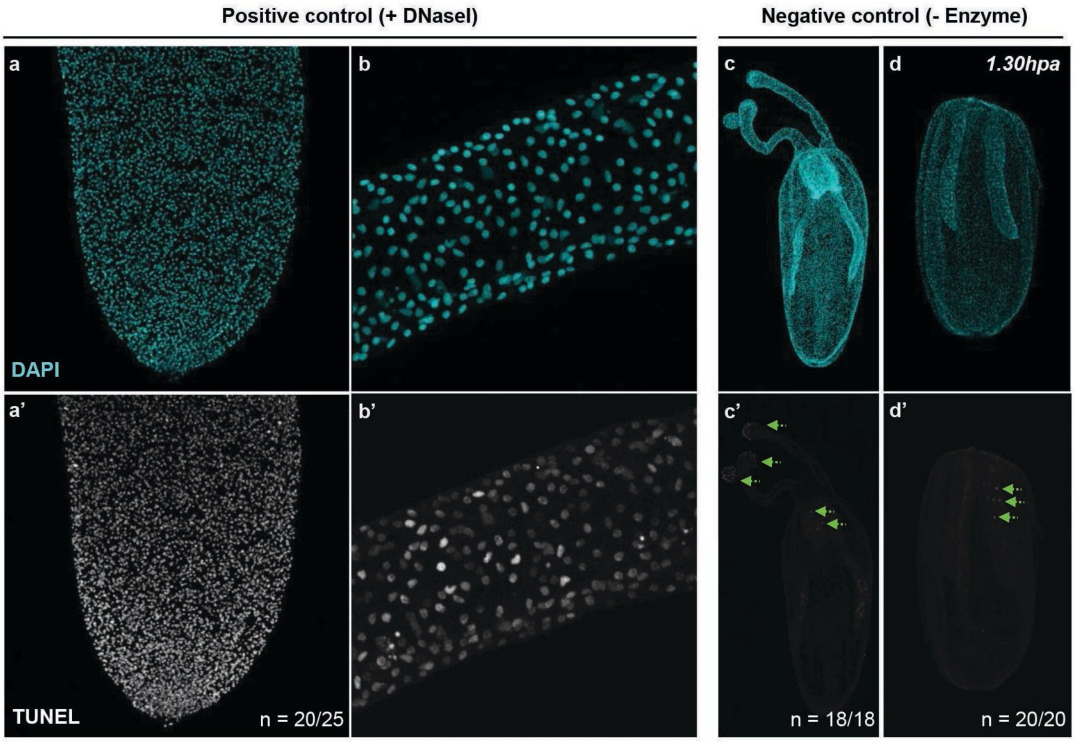

### Suppl Figure 4

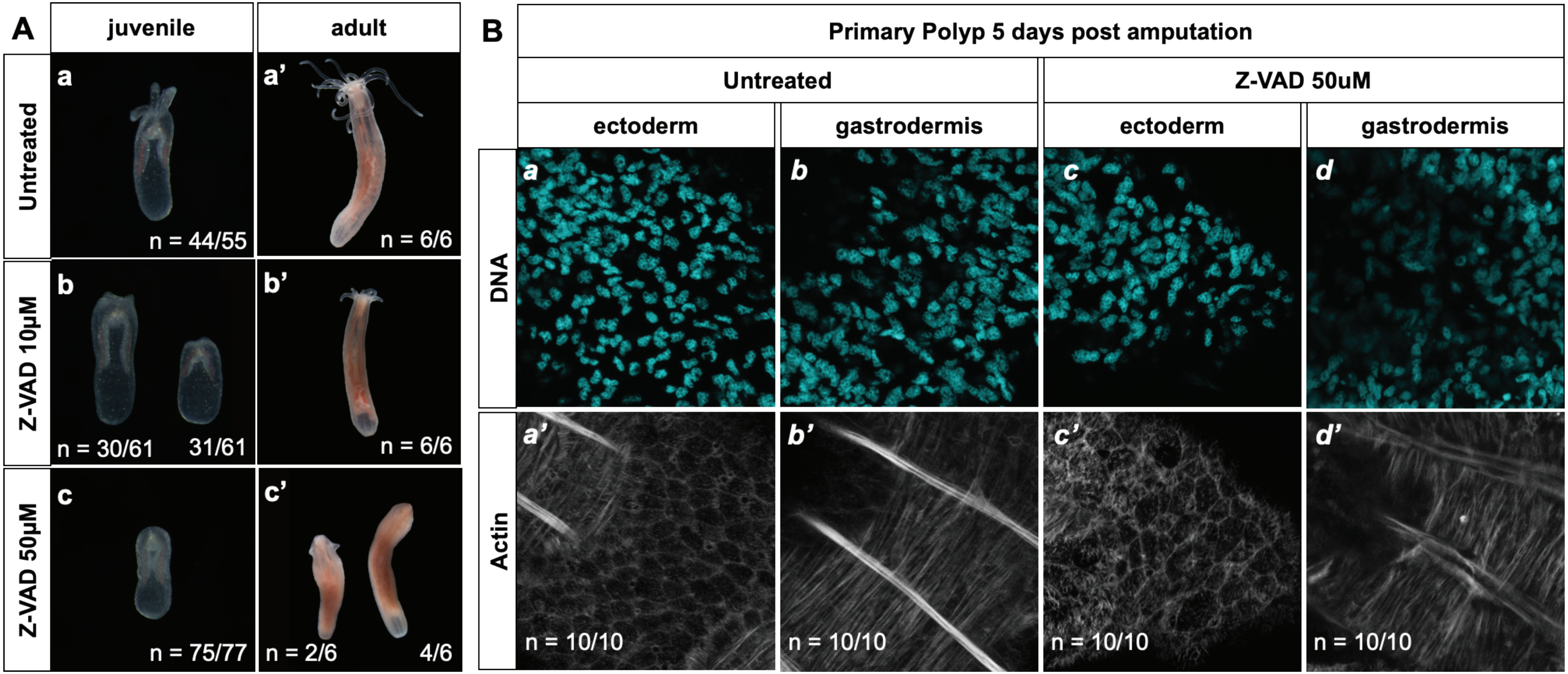

### Suppl Figure 5

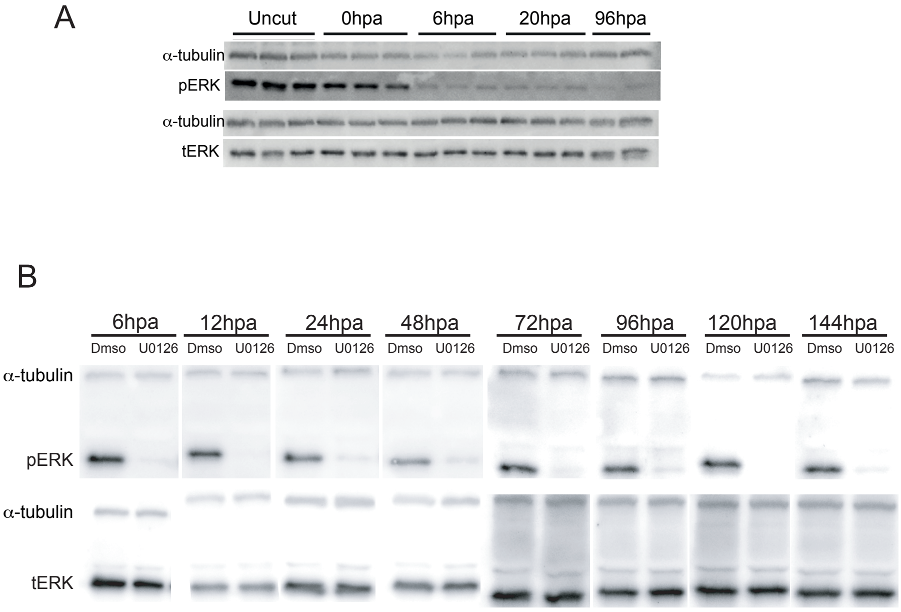

### Suppl Figure 6

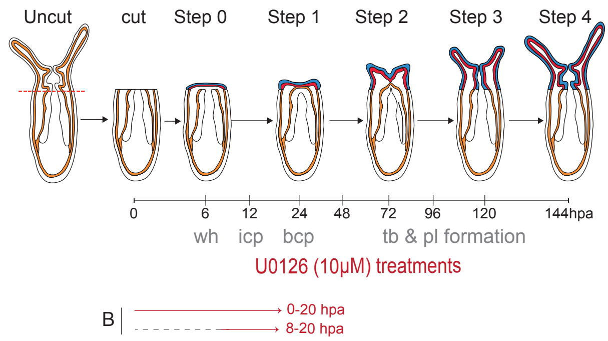

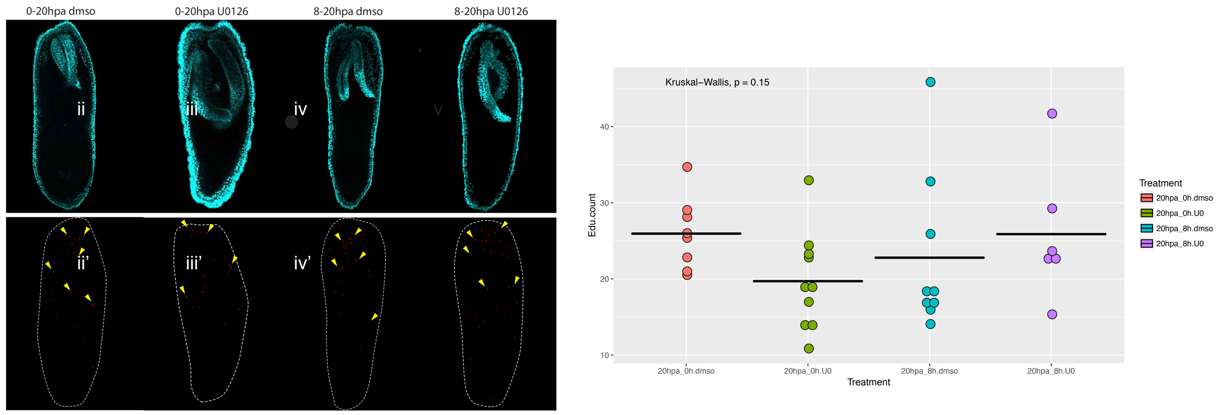

### Suppl Figure 7

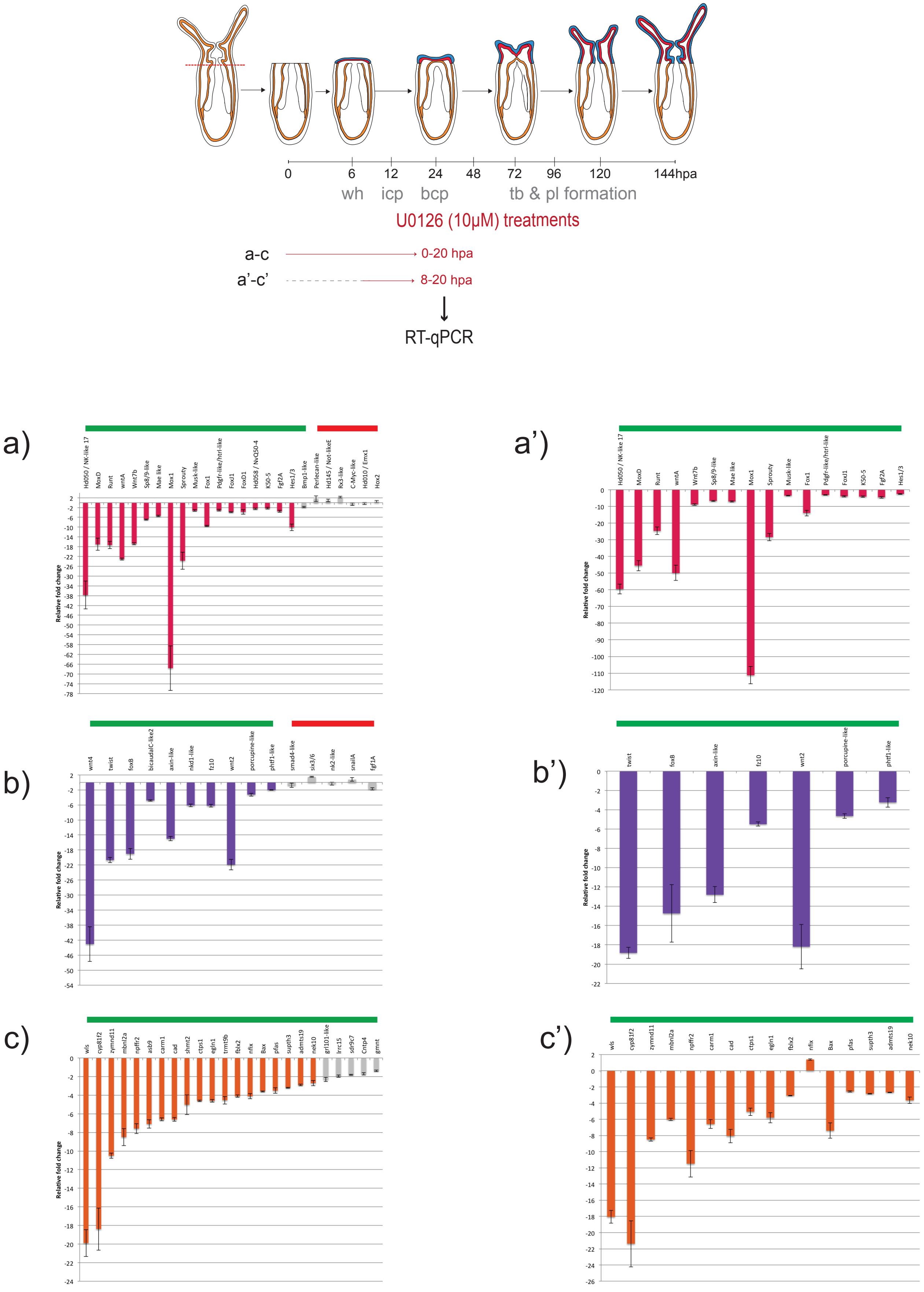
